# Experimental evolution of cellular miniaturization reveals a putative mechanism for cell size evolution

**DOI:** 10.64898/2025.12.06.692745

**Authors:** Garoña Ana, Lemos Victoria Morgane, Giometto Andrea, Fumasoni Marco

**Affiliations:** Gulbenkian Institute for Molecular Medicine (GIMM), Oeiras, Portugal; Cornell University, School of Civil and Environmental Engineering, Ithaca, USA

## Abstract

Cell volume, a key determinant of physiology, is maintained by cell size homeostasis. Large deviations from typical size are often harmful, yet cell sizes have diverged drastically in evolution. How size homeostasis evolves to support such diversity without impairing cellular function remains unclear. To address this question, we used experimental evolution to select progressively smaller *Saccharomyces cerevisiae* cells. Over 1,500 generations, we achieved a four-fold modal volume reduction compared to wild type. Despite this reduction, evolved populations maintained robust size homeostasis and relatively high fitness, revealing that substantial decreases in cell volume can occur without severe growth defects. Whole-genome sequencing and genetic perturbations identify mutations in the G1 cyclin Cln3 and the TOR pathway effector Sch9 as major contributors to cell size reduction, with the Greatwall kinase Rim15 signaling cascade further modulating this phenotype. Manipulating these pathways produced a six-fold range in cell size without changes in ploidy, with engineered mutations recapitulating the volumes of evolved lines and loss-of-function mutations yielding enlarged cells. Our results demonstrate the evolutionary plasticity of cell size homeostasis and reveal a putative mechanism for eukaryotic cell size evolution.

## INTRODUCTION

Cells are the fundamental unit of life, and their size is a key determinant of cell and organismal physiology (1). In microbial systems, variation in cell size can shape nutrient acquisition, metabolic scaling, and growth rate, making it a key target of evolutionary and ecological pressures (2). Through its effects on biosynthesis and proliferation, cell size constrains tissue architecture and regulates functional specialization (3–5).

Not surprisingly, cell size varies greatly across the tree of life. Among prokaryotes, cell diameters range from ∼0.2 μm in *Mycoplasma* species (6) to nearly 1 cm for *Thiomargarita magnifica* (7). Eukaryotes exhibit a similarly broad distribution, ranging from the smallest known free-living species *Ostreococcus tauri,* with a 1–2 μm diameter (8), to 15 cm long ostrich eggs (9). Even within multicellular organisms, such as humans, cells range from micron-scale lymphocytes to meter-long neurons (10).

Despite this large variability, specific cell types typically maintain their size within a narrow range through a process known as cell size homeostasis (11, 12). At its core, cell size homeostasis relies on the precise coordination of cell growth and division. The most direct evidence of such control on cell size is that cells born small typically grow more before dividing, and vice versa (13). While size control could in theory be performed throughout the cell cycle (14), most organisms exert it mainly prior to DNA synthesis, such as in budding yeast (15) and humans (16), or before nuclear division, such as in fission yeast (17). The general principles followed by cells to regulate their size, as well as the molecular mechanisms that execute this control, have been studied and debated for decades. Cell size control is best understood in the budding yeast *S. cerevisiae*, where cells are thought to commit to division once they reach a critical size (15) or after adding a constant volume each cycle (18). The mechanism that determines the transition from G1 to S phase has been proposed to depend mainly on titration of cell cycle activators (e.g., Cln3), inhibitors (e.g., Whi5), transcription factors (e.g., SBF), or their combination (for detailed mechanisms, see refs. (19–26)).

Strong evidence for the importance of cell size homeostasis is represented by the severe defects experienced by cells as they leave their narrow range of stereotypical sizes. Excessive cell size has been shown to cause cytoplasm dilution and genetic instability, ultimately increasing cells’ propensity to senescence (27–34). On the other hand, mutations causing the largest decrease in cell size impair the Target of Rapamycin (TOR) signaling and ribosome biogenesis, imposing severe reductions in cellular biosynthesis and fitness (35–38). Notably, altered cell size control has been observed in cancers (39–42).

It is therefore unclear how evolution has achieved changes in cell size of many orders of magnitude, when even slight increases or decreases are associated with severe cellular defects. These detrimental consequences could severely constrain the evolution of the cell size trait, posing an apparent paradox between the broad evolutionary diversity of cell sizes and the limited tolerance observed within species. To address this paradox, we sought to observe in real time the evolution of a large decrease in cell size in the model eukaryote *S. cerevisiae.* To this goal, we resorted to an experimental evolution regimen (43–45), simultaneously imposing direct selection on cellular size by Fluorescent-Activated Cell Sorting (FACS), followed by selection for competitive fitness. Applying this regimen over 1,500 generations produced miniaturized cells with modal volumes reduced four-fold relative to wild type (WT), with only marginal fitness tradeoffs. Miniaturized cells maintained their phenotype in the absence of selection and had size homeostasis comparable to their ancestor. Whole-genome sequencing revealed adaptive mutations in key players of the G1 cyclins and TOR pathway, with additional contributions from components of the Greatwall kinase cascade. Reconstruction of these mutations showed that their combined effects generate markedly smaller cells while maintaining near-wild-type fitness, thereby decoupling cell size from growth rate. Targeted engineering of these pathways substantially altered cell size control and cell cycle dynamics, yielding a six-fold range in cell volume. Our work demonstrates the plasticity of the mechanisms governing cell size homeostasis and suggests a putative mechanism for cell size evolution.

## RESULTS

To investigate how cell size can evolve under sustained proliferative capacity, we developed an experimental system in *S. cerevisiae* that alternates two selective regimens targeting cell size and reproductive fitness, respectively. Cell populations were analyzed by FACS, using the light scattered by cells (forward scatter area, FSC-A), a widely used proxy for their size (46). Only the smallest 7% of cells were collected into separate tubes and subjected to the second selection step (Figure 1A; see SI Appendix and Figure S1A for details). The second selection consisted of the unconstrained growth of the sorted population for at least 11 generations, during which cells competed for resources (Figure 1A). After reaching marked turbidity, typical of cells reaching stationary phase, the three parallel small (S1-3) populations were subjected again to both selection regimens. Repeating this cycle daily resulted in approximately 1,500 generations of evolution (Figure 1B). Under these conditions, selection favors mutations that reduce cell size without compromising proliferation. To speed the evolution, the starting haploid population (ancestor) carried a mutator allele in the catalytic subunit of DNA polymerase delta (*pol3-L523D* (47), Table S1). In addition, we engineered the ancestor strain to express a cytoplasmic fluorescent protein (ymCitrine) to prevent subcellular debris from being sorted.

**Figure 1.**
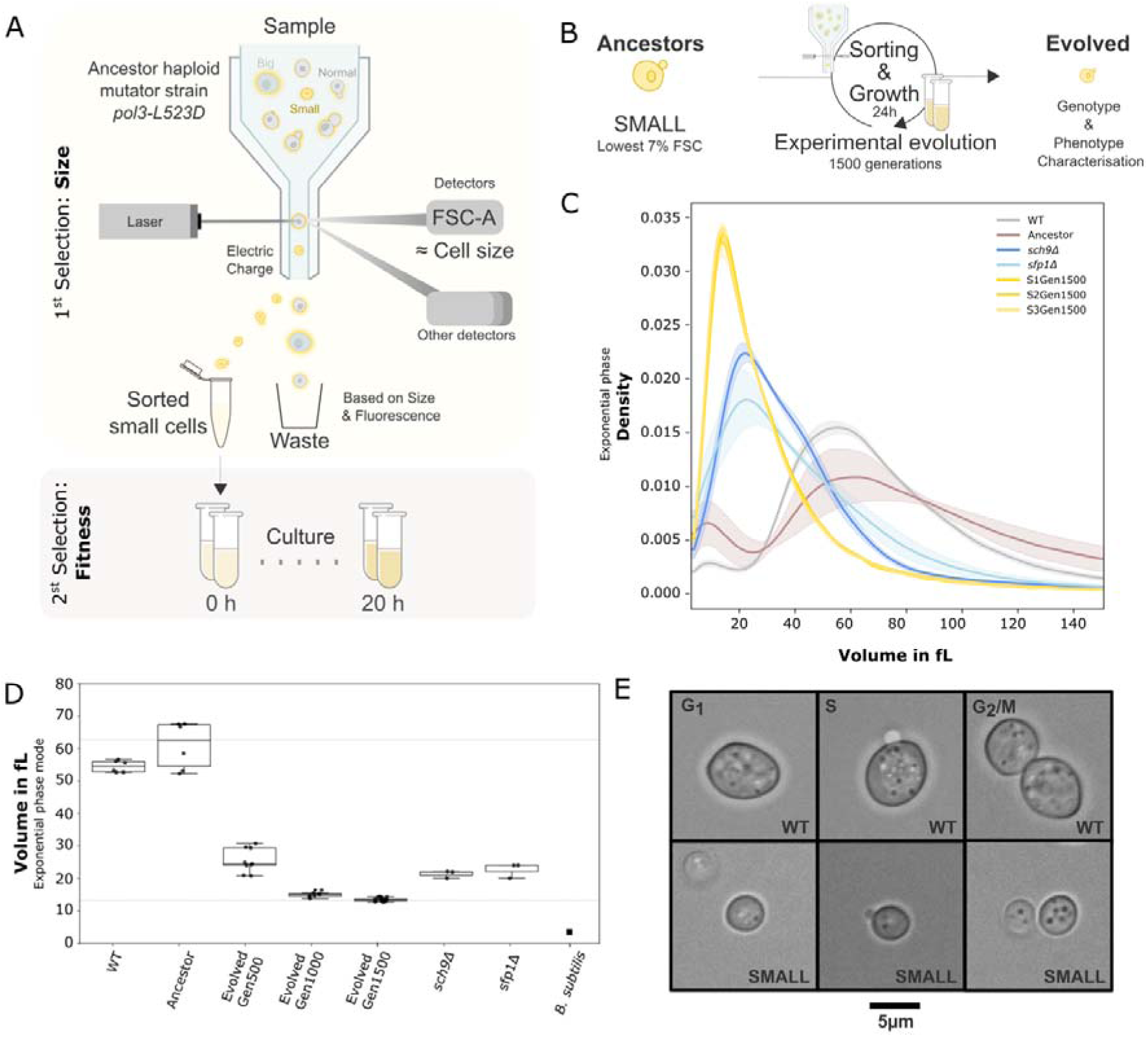
Experimental evolution of cellular miniaturization. (A and B) Schematic representation of size selection and evolution regimens. A haploid *pol3-L523D* mutator ancestor population was sorted into three small (S) subpopulations and subjected to daily size selection. Cells in the lowest 7% of the forward-scatter distribution were sorted (100,000 cells) and cultured in 10 mL rich medium (YPD). Populations were serially grown and sorted for 116 cycles (∼1,500 generations), with samples collected and stored every 24 h. (C) Cell volume distributions of exponential-phase cultures. Smooth density curves represent kernel density estimates fitted to histograms of cell volumes measured with a Coulter counter. For each strain, densities were computed per replicate and averaged to obtain a mean density curve. Shaded areas represent the standard deviation across replicates. In large cell size strains, the first peak corresponds to debris rather than intact cells, a known Coulter counter artifact that is captured as an artificial peak in the density estimate. (D) Modal cell volume from measurements of exponential-phase cultures (Coulter counter) across generations. Data from the three S-populations are plotted together (“evolved”), showing progressive evolution toward smaller size, reaching up to a four-fold reduction in volume. Grey lines indicate wild-type and final evolved sizes. Box plots show medians, interquartile ranges, whiskers (1.5× IQR). (E) Light micrographs of wild-type and evolved small-cell populations at different cell-cycle stages (G1, S, G2/M). Scale bar, 5 µm.

Over 1,500 generations, the evolving populations (S1–3) showed a gradual decrease in FCS values as measured by flow cytometry (Figure S1B). Microscopy and Coulter counter analyses confirmed that this reduction was accompanied by a marked decrease in cell volume (Figure 1C–E). Literature values from Coulter counter measurements are typically reported from exponential-phase cultures, when cells divide rapidly in the presence of abundant nutrients. In contrast, in our experimental protocol, size selection occurred as cultures approached the stationary phase, when nutrients become limiting and cell division progressively slows. To determine whether the evolved reduction in cell size was maintained across growth phases, we measured cell volumes in both exponentially growing and saturated cultures (Figure 1C-D and Figure S1C–E). In exponential phase, WT and Ancestor cells exhibited unimodal size distributions with modes of 54.7 ± 1.6 fL (mean ± SD) and 61.2 ± 7.29 fL (mean ± SD), respectively. By generation 1,500, the S-populations displayed an approximately four-fold reduction in modal size over WT (12.8–13.8 fL; Figure 1C-D), accompanied by ∼2.2-fold and ∼2.7-fold decreases in mean and median volumes, respectively (means ∼30–31 fL; medians ∼22–23 fL), approaching volumes closer to those of the bacterium *B. subtilis* than to their own ancestor. Comparatively, perturbations of the TOR signaling pathway (*sch9*Δ and *sfp1*Δ), which produce some of the smallest previously described yeast cells (35, 48), only reduced modal volumes by ∼2.5-fold (22.8 ± 2.9 fL and 22.6 ± 1.9 fL mean ± SD, respectively, Figure 1C). Upon entry into stationary phase, WT populations displayed a broader, and frequently bimodal volume distribution, with peaks corresponding to small daughter cells (∼22 fL) and larger mother cells (∼80 fL) (Figure S1C-D). In contrast, the evolved populations remained unimodal, with a single peak around ∼11 fL. Because the WT stationary-phase distribution reflects a mixture of cell types, we compared median cell volumes, which revealed that the evolved lines remained approximately 2.5-fold smaller than WT cells, indicating a robust size reduction also at the end of the growth curve (Figure S1E). Unless otherwise noted, all cell size measurements reported below were obtained from exponentially growing cultures.

Together, these results indicate that, to the best of our knowledge, this selection experiment produced the smallest *S. cerevisiae* cells reported to date under standard laboratory growth conditions. Modal cell sizes as low as 16 fL have previously been reported for mutants affecting the G1/S transition when grown in poor carbon sources such as glycerol; however, these conditions markedly slow cell growth and division (20, 35, 49, 50) (Figure S1F). Comparable size reductions in *S. pombe* have been reported to cause mitotic catastrophe (51, 52), highlighting the severe constraints on extreme size reduction. Strikingly, the evolved cells reached volumes similar to those of the smallest species within the *Saccharomycotina* subphylum (Figure S1G), spanning nearly half of the natural size diversity observed across the phylum, despite over 400 million years of divergence (53).

### Gradual cell size decrease reshapes cell cycle dynamics, cellular morphology, and nutrient-dependent size control

The dramatic changes in cell volumes observed during our evolution experiment prompted us to examine potential alterations in cellular architecture. The cell cycle is a major determinant of cell morphology in budding yeast (54). To determine whether changes in cell size were associated with alterations in the cell cycle, we analyzed DNA content in exponentially growing populations, where signal intensity distinguishes cell cycle phases (1C for G1, 2C for G2/M, and 1C < DNA < 2C for S phase). Strikingly, by generation 1,500, evolved small cells displayed a near absence of G1 compared with WT (Figure 2A and S2A). Notably, cell cycle changes during evolution occurred in two steps: during the first ∼150 generations, the proportion of cells in G1 increased, followed by a sharp shift around generation 700 that resulted in the near disappearance of G1 (Figure 2B and S2B). Calculating the absolute time spent in each phase confirmed this trend, showing that the reduction of G1 duration was primarily compensated by a corresponding extension of S/G2/M phases (Figure 2B-C and S2B).

**Figure 2.**
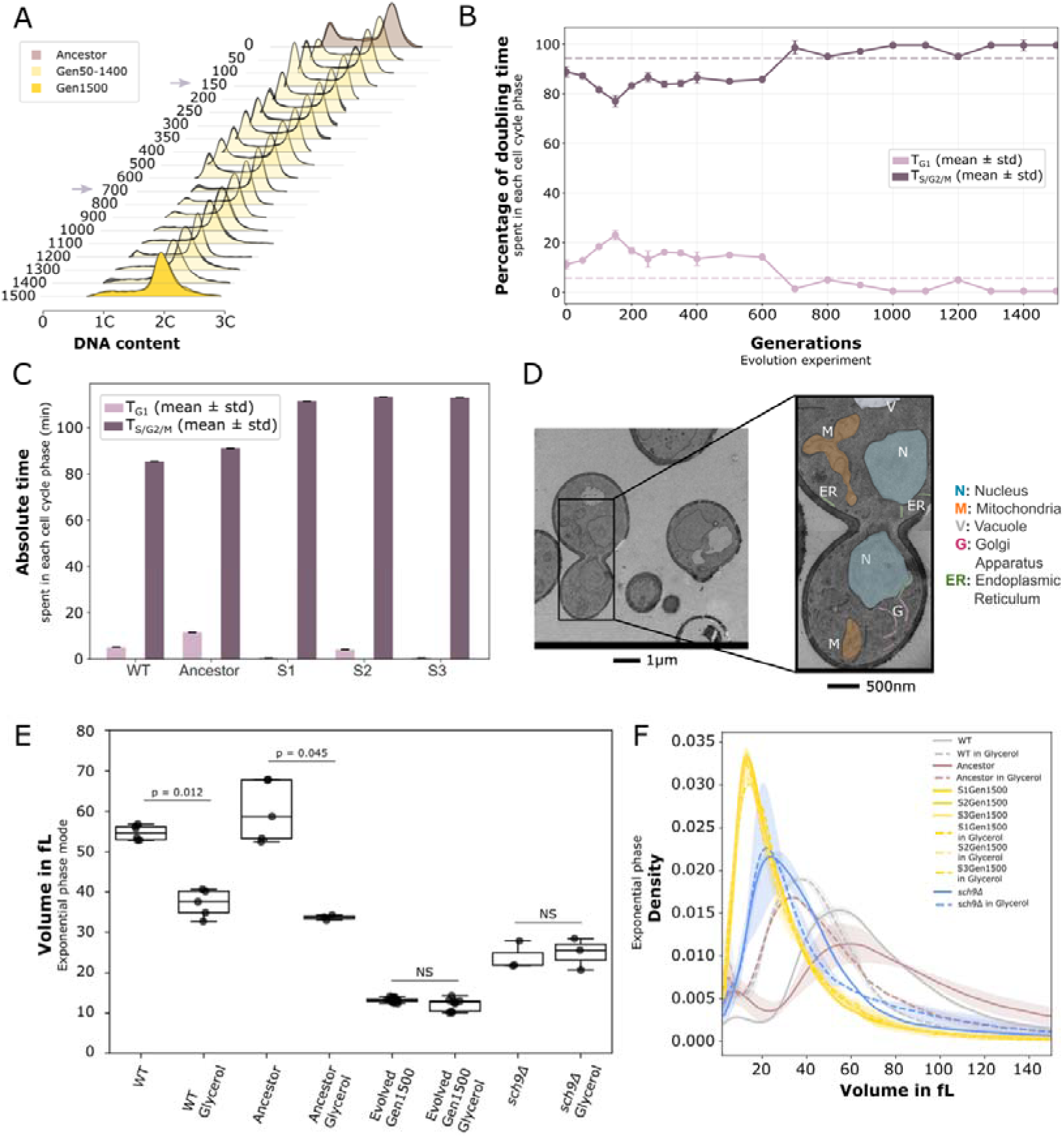
Physiology of cell miniaturization. (A) DNA-content profiles of ancestral and representative evolved (S1) populations during exponential growth across 1,500 generations, measured by flow cytometry. Lines show means; shaded areas, SD (n = 3). G2 peak height was normalized to 1. Arrows indicate time points of cell cycle profile changes (see main text). (B) Percentage of doubling time spent in G1 and S/G2/M phases for the S1 population at generation 1,500. Bars show mean ± SD (n = 3). (C) Absolute time spent in each cell cycle phase, calculated as the fraction of the doubling time allocated to each phase (56). (D) Transmission electron micrograph of evolved *S. cerevisiae* cells during exponential growth in YPD. (E) Modal cell volume measurements (Coulter counter) grown in glucose versus glycerol as carbon sources. Data from the three S-populations are plotted together (“evolved”). Box plots show medians, interquartile ranges, and whiskers (1.5× IQR). Pairwise comparisons between carbon sources (glucose vs glycerol) were performed using the Mann–Whitney U test; p-values are: WT (p = 0.012), Ancestor (p = 0.045), evolved-Gen1500 (p = 0.051), and *sch9*Δ (p = 1.000). (F) Cell volume distributions of exponential-phase cultures grown in glucose or glycerol as a carbon source. Smooth density curves represent kernel density estimates fitted to histograms of cell volumes measured with a Coulter counter. For each strain, densities were computed per replicate and averaged to obtain a mean density curve. Shaded areas represent the standard deviation across replicates.

We next focused on intracellular organization, reasoning that reduced volume might occur at the expense of specific organelles. To obtain a general view of subcellular structures, we performed transmission electron microscopy (55) on exponentially growing WT, ancestral, and evolved S-populations. Buds smaller than mother cells, consistent with asymmetric cell division (15), were commonly observed. Interestingly, major organelles displayed their typical morphology, though with altered proportions, suggesting a potential differential miniaturization of organelles (Figure 2D and Figure S2C). Despite the presence of petite clones (cells lacking functional mitochondrial respiration) within the populations, most evolved isolates, including the smallest, retained growth on non-fermentable carbon sources, indicating preserved mitochondrial function. Petite clones did not differ in cell size from non-petites (Figure S2D).

Given preserved respiratory capacity, we tested whether evolved cells maintain the canonical relationship between nutrient quality and cell size. Typically, cells decrease in volume in lower-quality media (e.g., non-fermentable carbon sources) relative to high-quality media such as glucose. When cultured in the non-fermentable carbon source glycerol, WT and ancestor cells reduced their modal cell size from 55 and 60 fL to 37 and 34 fL, respectively, corresponding to a ∼1.5–2-fold decrease in modal cell volume. In contrast, evolved populations maintained almost identical modal cell sizes in glucose and glycerol (13.22 vs. 12.12 fL; Figure 2E-F and Figure S2E). This result suggests either that evolved cells have reached a minimal attainable cell volume, or that the molecular pathway linking nutrient quality to cell size has been altered to mimic poor-nutrient conditions even in high-quality media like glucose.

Together, these observations indicate that cellular miniaturization in evolved populations is accompanied by extensive remodeling of cell cycle progression, cellular organization, and nutrient-dependent size control.

### Evolved cells decouple miniaturized size from pronounced slow growth

Marked decreases in cell size, whether caused by mutations, poor nutrients, or a combination of both, are typically associated with pronounced slow growth. We therefore wondered how changes in cell size affected the evolved populations’ growth rates. Growth curves of WT, ancestral, and S-populations revealed a marked increase in maximum population density in the evolved lines (Figure 3A). Unexpectedly, their growth rates were only marginally reduced relative to ancestors, which themselves grew slower than WT, potentially due to the mutator phenotype (Figure 3B and Figure S3A-C). To precisely estimate this defect, we resorted to pairwise competition assays, which measure relative fitness, a measure of reproductive success of organisms (57). All three evolved S-populations had equivalent relative fitness (–7,79% ± 1.4, mean ± SD vs. Ancestor), whereas WT showed a ∼5% advantage relative to the ancestor (Figure 3C). In contrast, deletion of *SCH9*, which produces some of the smallest cells in rich media, caused a severe fitness defect (∼24% vs. Ancestor & ∼28% vs. WT) despite reducing modal cell volume only 2.5-fold in exponential phase. Across all populations, fitness followed a non-uniform trajectory: (i) declining during the first 200 generations, (ii) recovering by generation ∼250, (iii) dropping sharply until ∼500, and (iv) stabilizing with minor decreases until 1,500 (Figure 3D, S3D-G). Fitness was more strongly associated with cell-cycle dynamics than with cell volume (Pearson r = 0.67, p < 0.001 for G1 fraction; r = 0.30, p = 0.11 for cell volume), indicating that changes in fitness primarily reflect alterations in cell-cycle progression (Figure 2B and 3D). Importantly, these results highlight how evolved lines fall outside of an inverse linear relationship between cell size and doubling time observable across a wide range of mutants (Figure 3E).

**Figure 3.**
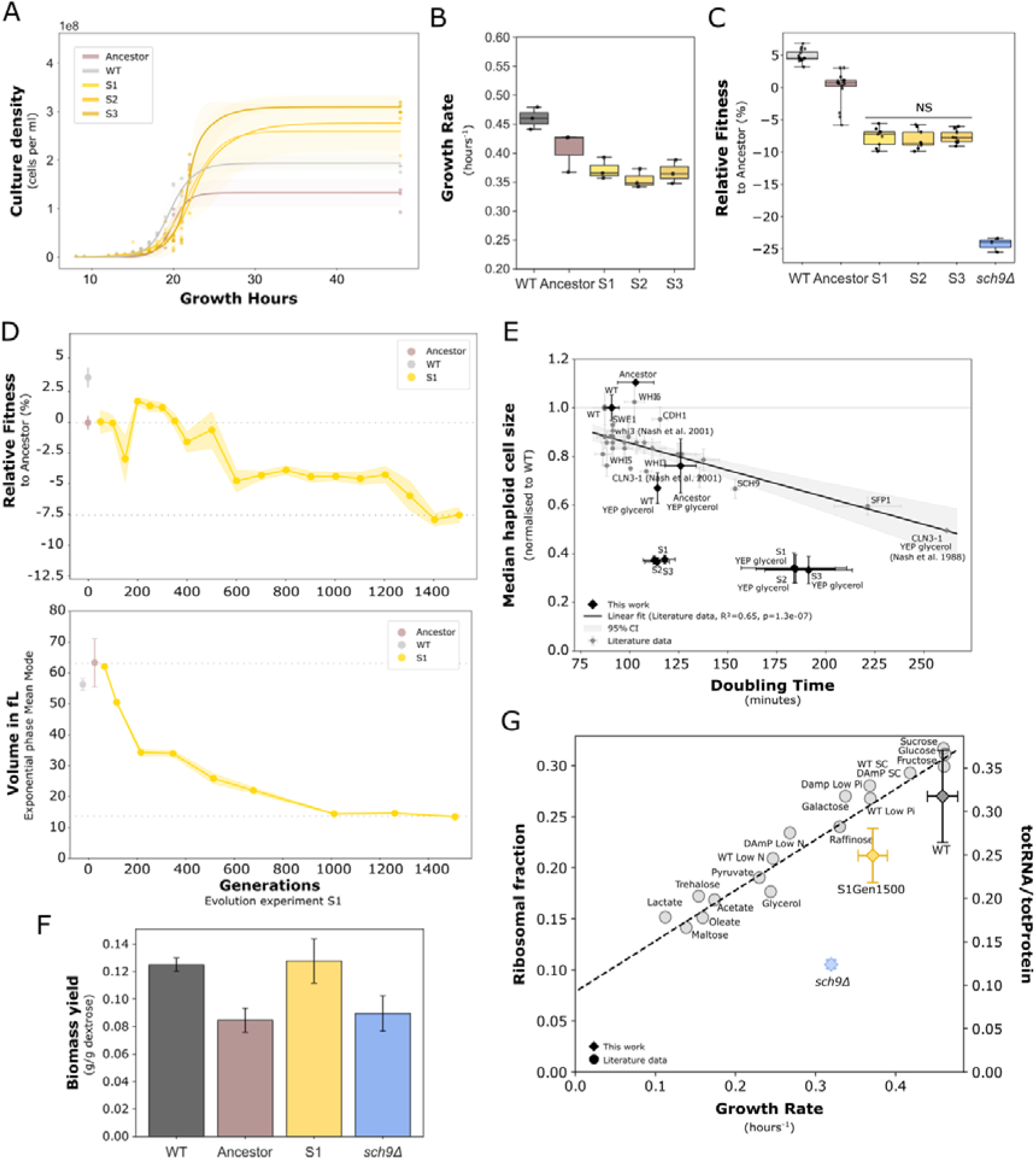
Fitness and biomass production. (A) Growth curves of selected strains measured as cell density (cells/mL) versus time (h). (B) Maximum growth rates of reference and evolved strains (generation 1,500) in YPD. Box plots show medians, interquartile ranges, and whiskers (1.5× IQR). (C) Relative fitness of wild-type, ancestral, and evolved populations (S1–S3) at generation 1,500. Relative fitness is the ratio of reproductive success of a strain to that of a reference strain under identical conditions. Fitness of evolved and *sch9*Δ strains differed significantly from the ancestor and wild type (adjusted p < 0.01), with no significant differences among evolved lines. Box plots as in (B). Full pairwise statistics are provided in Supplementary file: RawData-Figures. (D) Fitness and cell-size trajectories across 1,500 generations for the S1 population. Lines show mean values; shaded areas show SD. (E) Relationship between fold decrease relative to WT median cell size and doubling time in *S. cerevisiae*. Circles represent data from (35, 49, 50). Diamonds represent strains measured in this study. Error bars represent propagated measurement uncertainty. The solid black line shows a linear regression computed using only mutant strains, with the shaded area representing the 95% confidence interval. Genes of interest are annotated. (F) Biomass derived from dry mass per cell, converted to yield relative to dextrose content in the medium. (G) Scaling of the ribosomal fraction of the proteome with the population’s growth rate. Data from (60) (circles) are compared with experimental measurements obtained in this study (diamonds). A linear regression (dashed line) was fitted to the ribosomal fraction values in (60). The total RNA-to-protein ratio (RNA/protein) is plotted on a secondary y-axis as a proxy for the ribosomal fraction of the proteome.

The most pronounced decreases in cell size have previously been associated with nutrient limitation or mutations in nutrient-sensing pathways. These conditions were proposed to modulate cell size through reduced ribosome biogenesis (48). However, numerous microbial physiology studies have linked reduced ribosome biogenesis with decreased growth rates (58, 59). This raises the question: how do evolved cells maintain a miniaturized cell size while sustaining growth rate and fitness? To address this, we examined the biosynthetic capacity of the evolved lines. At saturation, the dry mass produced per gram of dextrose by evolved lines was comparable to that of WT, whereas *sch9*Δ cells produced approximately one-third less (Figure 3F). We next estimated the ribosomal mass fraction of the proteome from the RNA/protein ratio, a commonly used proxy for ribosome biogenesis in microbial physiology, also applicable to yeast (59). As expected, *sch9*Δ cells showed a marked reduction in ribosomal mass fraction, whereas the evolved lines maintained ribosome biogenesis at levels closer to WT (Figure 3G).

Altogether, these results suggest that evolved lines maintain efficient growth despite a markedly reduced cell volume, by preserving ribosome biogenesis and biomass production.

### New evolved size homeostasis is stable and robust

How stable is the evolved miniaturized size? Measurements performed right after the last sorting round may reflect a transient physiological state, rather than an evolutionarily stable phenotype. To address this possibility, clones were isolated from two independent populations (S1Cl2 and S2Cl4) and passaged for 150 generations in the absence of any size selection. Interestingly, S-clones stably maintained their size across the experiment, resulting in a final fold change similar to that observed in control WT lines (Figure 4A and Figure S4A-D). In contrast, *sch9*Δ lines consistently reverted to WT sizes between generation 50 to 100 despite starting with a larger volume than S-clones (Figure 4A and Figure S4A-D). A key question is whether miniaturized cells actively regulate their size, or whether they merely preserve the volumes attained at the end of the experiment. To address this, we transiently arrested cells at the G1/S transition using the pheromone α-factor, which permits growth while blocking division (61) (Figure S4E-F). While the arrest was not fully efficient in S-clones, it nevertheless led to a ∼1.8-fold increase in cell volume in both WT and evolved cells. The increased volume, however, was progressively reversed upon release into fresh medium, indicating that cells actively return to their characteristic small size (Figure S4E, S4G).

**Figure 4.**
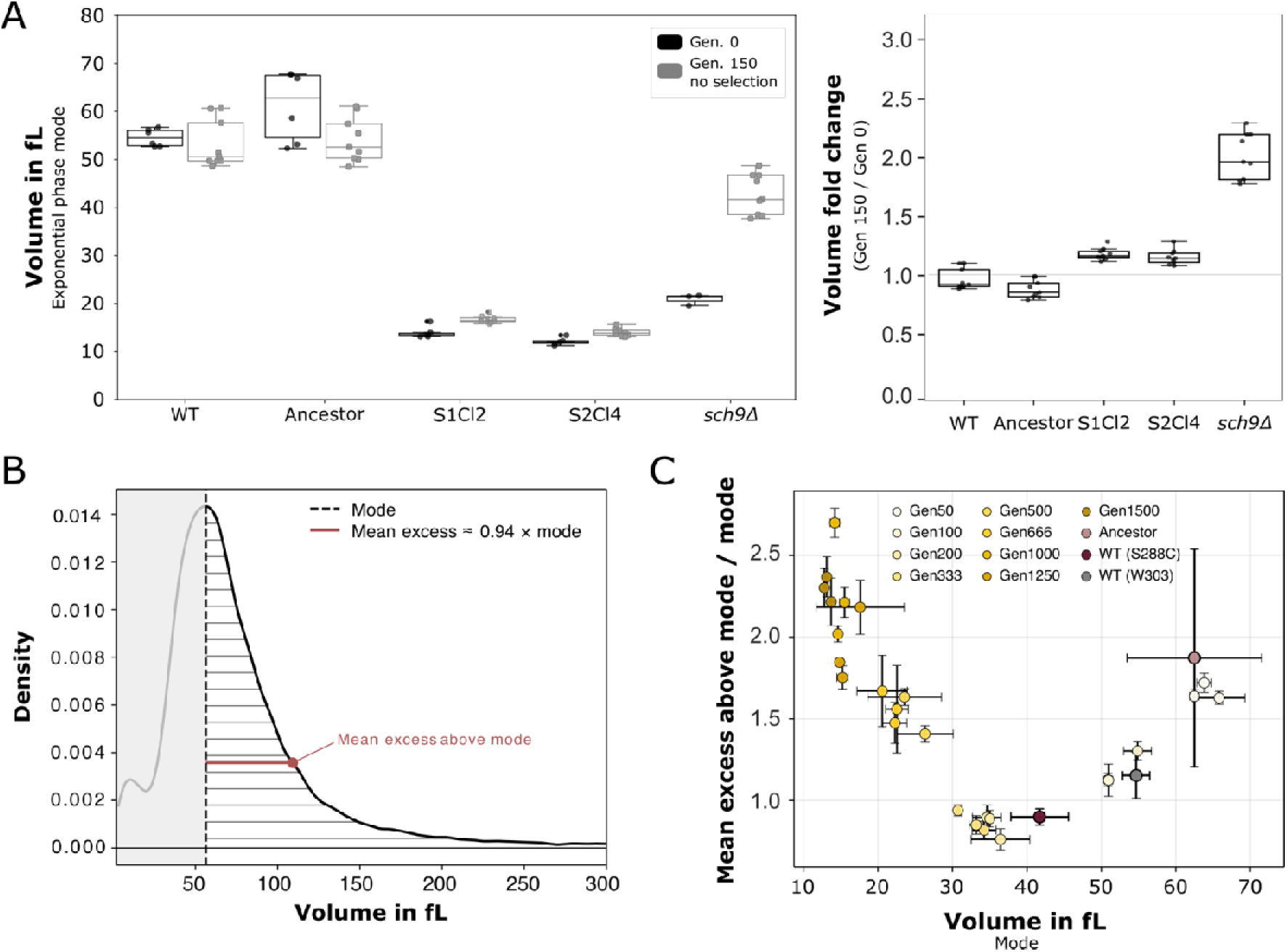
Evolution of cell size homeostasis. (A) Stability of cell size during unconstrained experimental evolution. Left: cell-size measurements of WT, Ancestor, *sch9*Δ, and evolved clones at 1,500 generations from populations evolved for 150 generations without size selection. Box plots show medians, interquartile ranges, and whiskers (1.5× IQR). Pairwise comparisons between strains were performed using the Mann–Whitney U test: WT vs WTGen150 (p = 0.328), Ancestor vs AncestorGen150 (p = 0.082), S1Cl2 vs S1Cl2Gen150 (p = 0.011), S2Cl4 vs S2Cl4Gen150 (p = 0.008) and *sch9*Δ vs *sch9*ΔGen150 (p = 0.015). Right: fold change in cell size relative to time 0. (B) Our metric of cell size distribution width calculates the mean excess above the mode, which is a generalization of the half-width at half maximum that accounts for the distribution width across all density values, rather than just the half maximum (SI Appendix). We focus on the right tail of the distribution to avoid spurious effects due to the presence of cell debris at small cell volumes, highlighted here by logarithmic cell volume binning. Shown here is the distribution of WT (W303). (C) Width of cell size distributions, quantified as the mean excess above the mode (SI Appendix), normalized by modal volume. The Ancestor (blue, ∼62.5 fL modal volume, normalized mean excess ∼2.8) had a broad right tail relative to WT, leading to a large normalized mean excess. Evolved populations progressively reduced both modal volume and normalized mean excess over generations, with early-generation populations (Gen100–Gen200) converging near WT (W303) values (∼50-55 fL, normalized mean excess ∼1.2). At modal volumes below ∼30 fL, the normalized mean excess rose again to ∼2.5, approaching the ancestral value.

The ability of miniaturized cells to maintain their size across many generations and under transient perturbations reflects efficient size homeostasis, which is expected to manifest as a narrow distribution of cell sizes around the population mean. To test this prediction, we compared the width of cell size distributions across WT, ancestral strains, and evolved populations at successive generations (Fig. 4A). To avoid spurious contributions from cell debris and non-cellular particles at low volumes, which bias conventional statistics such as the mean and standard deviation, we quantified distribution width as the mean excess above the modal volume E[v - v_mode_|v > v_mode_], normalized by the modal volume (SI Appendix). This dimensionless metric captures the average relative width of the right tail of the size distribution independently of debris at low volumes (Figure 4B). Over the course of evolution, populations initially reduced both their modal volume and their normalized mean excess, converging toward values below those of the ancestor and comparable to WT (Figure 4C). Once modal volumes fell below ∼30 fL, however, normalized mean excess increased again, returning to values comparable to the ancestor despite a dramatically reduced characteristic cell size. Overall, miniaturized S-clones thus maintain a degree of size homeostasis similar to the ancestor, demonstrating that the reduction in cell size was not accompanied by a proportional loss of size control.

Together, these results demonstrate that miniaturized cells have evolved a new cell size homeostasis that is evolutionarily stable, robust to transient perturbations, and capable of maintaining population-level size control as tightly as their own ancestor.

### Genetic landscape of cellular miniaturization

We sought to determine the genetic basis of the evolved phenotype. Evolved cells retained their characteristic size when co-cultured with wild-type strains, indicating that miniaturization was not driven by secreted or diffusible factors (Figure S5A). The phenotype’s persistence over 150 generations in the absence of selection (Figure 4A) strongly suggested a genetic basis. Supporting this idea, crosses between evolved clones and WT produced diploids of intermediate size between haploid and diploid WT, consistent with the presence of both recessive and dominant mutations. Furthermore, crosses between evolved clones and mating-type–switched clones yielded diploids that retained a large reduction in cell volume relative to WT diploids (Figure S5B), suggesting that the reduced cell size is independent of ploidy.

To identify causative mutations, we performed whole-genome, whole-population sequencing across multiple evolutionary timepoints. As expected from the mutator allele present in the ancestral population, we detected 869 new unique mutations that arose during evolution in coding and regulatory regions across the three populations (Figure 5A, Table S2). By the end of the experiment, an average of 267 single-nucleotide polymorphisms (SNPs) and small insertions/deletions (indels) persisted per population (285 in S1, 301 in S2 and 216 in S3, Figure S5C). No aneuploidies, segmental amplifications, or large-scale changes in DNA copy number were observed, thus confirming the presence of a full haploid genome (Figure S5D). The evolved populations exhibited distinct clonal compositions (Figure S5E), however, several alleles were shared across populations. These mutations were absent from the ancestral strain and, in most cases, involved identical nucleotide substitutions. The probability that a large number of identical mutations arose independently across populations is therefore extremely low, pointing towards a likely cross-contamination during the FACS selection steps. Accordingly, the evolved lines were treated as a single evolving population for subsequent analyses rather than as independent replicates (Figure 5A).

**Figure 5.**
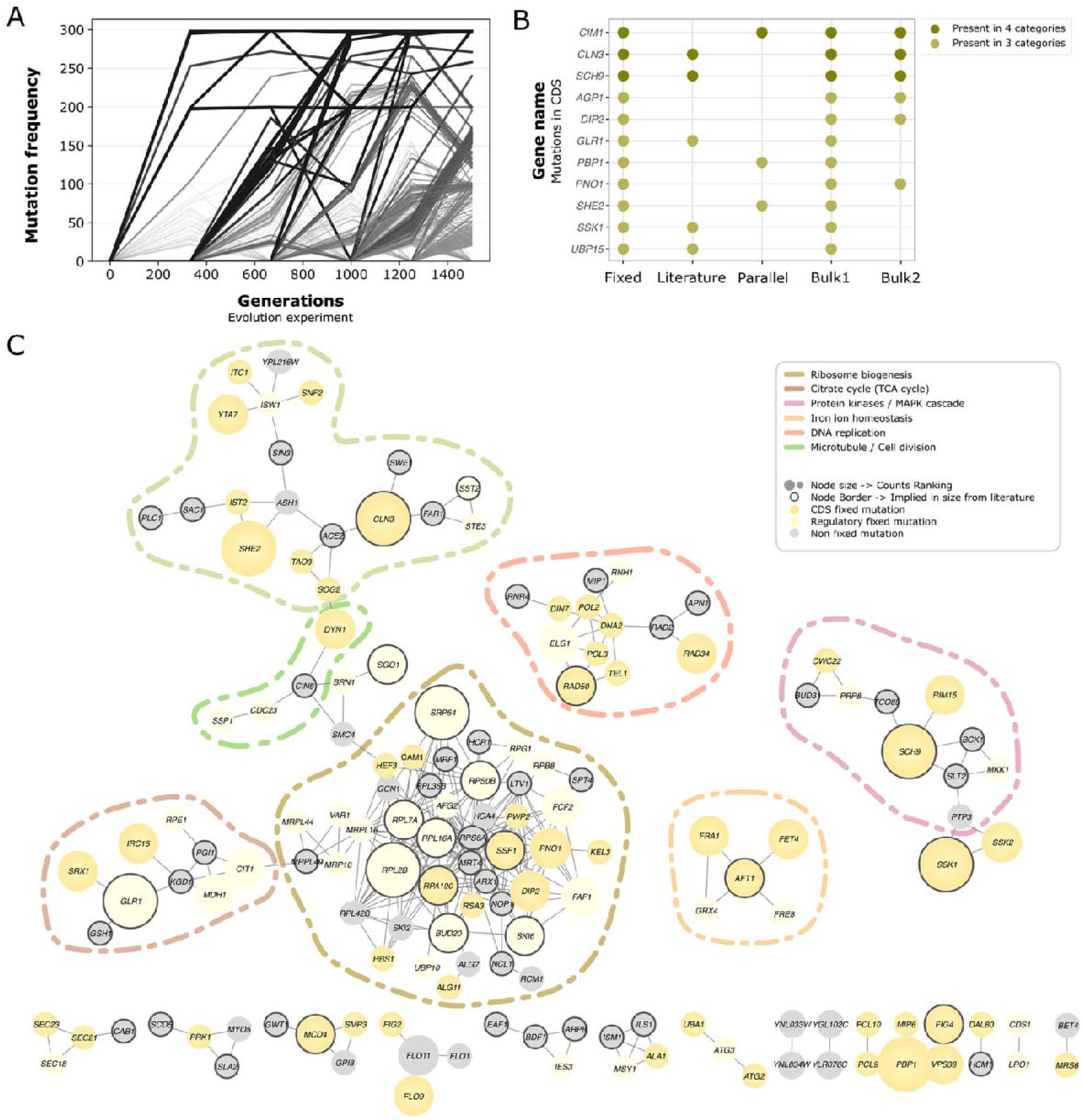
Genomic landscape of the evolved phenotype. (A) Allele-frequency trajectories of mutations detected in the combined population (S1–S3). A 300% frequency indicates fixation in all three populations. Line thickness reflects the final frequency of the mutation at generation 1,500. (B) List of putative adaptive gene mutations ranked by the number of satisfied criteria outlined in the main text and affected by non-synonymous mutations located within coding sequences (CDS). Only genes appearing in at least three categories are included (full list in Table S6). (C) Interaction network of mutations detected in S1–S3 evolved populations curated in Cytoscape (52). Grey lines indicate known genetic and physical interactions (STRING database (53)). Node diameter reflects the number of satisfied criteria we used to detect putative adaptive mutations. Only genes appearing in at least two categories are included. Nodes are color-coded: dark yellow for non-synonymous mutations within CDS fixed at generation 1,500, light yellow for mutations in the regulatory regions fixed at generation 1,500, and grey for non-fixed mutations. Bold outlines mark genes linked to cell size changes in the literature (either decrease or increase).

To identify putative adaptive mutations associated with the evolved cell size, we employed four non-exhaustive, but complementary approaches: (i) tracking lineages that rose to fixation in any of the three sub-populations, identifying cohorts of mutations under positive selection (Table S2); (ii) searching for genes previously implicated in cell size regulation (Table S3); (iii) using statistical inference to identify genes mutated more frequently than expected given their length and the observed mutation rates (parallel evolution; Table S4); and (iv) performing two consecutive rounds of backcrossing and selection for small cell size to pinpoint mutations segregating with the phenotype (bulk segregant analysis; Table S5; see S1 Appendix for details). A summary of the genes identified by these methods, as well as their reported interactions, is portrayed in Figure 5B-C (Table S6). Despite several genetic and physical interactions among the identified genes, no gene ontology term was found to be statistically enriched.

Our analysis indicates that the miniaturized phenotype is heritable and genetically encoded, but the underlying adaptive landscape is complex. The large number of candidate mutations likely includes false positives arising from hitchhiking or linkage, as well as variants under indirect selection. Some of these may reflect selection on traits influencing the flow-cytometry size proxy (FSC-A) rather than cell size itself, or compensatory adaptations to physiological changes associated with miniaturization (62).

### Combined mutations in the G1 cyclin and Greatwall signalling cascades cause large deviations in cell size

We set out to identify the mutations primarily responsible for the change in cell size among those identified as putatively adaptive. *CIM1*, *CLN3*, and *SCH9* ranked highest in our analysis and were therefore selected for further investigation. *CIM1* encodes for a mitochondrial HMG-box protein that limits mitochondrial DNA copy numbers (63), and independently accumulated up to six mutations in the protein over the course of the experiment, including three frameshifts resulting in premature stop codons (Figure S6A). Cln3, a G1 cyclin that partners with Cdk1 to drive the G1/S transition by activating phase-specific gene expression, has been shown to reduce cell size when truncated at its C-terminus (e.g. *cln3-1*), due to increased protein stability (49, 64, 65). Our evolved strains carried a similar truncation, *cln3-S410\**, just downstream of the *cln3-1* allele (Figure 6A). Sch9 is a key effector of the TORC1 pathway, which integrates nutrient sensing with anabolic and catabolic processes. We identified a mutation (*sch9*-L343S) within the conserved C2 domain (residues 184–402), a calcium-dependent membrane-targeting module that may also inhibit the kinase domain (residues 403–738; Figure 6A, (66)).

**Figure 6.**
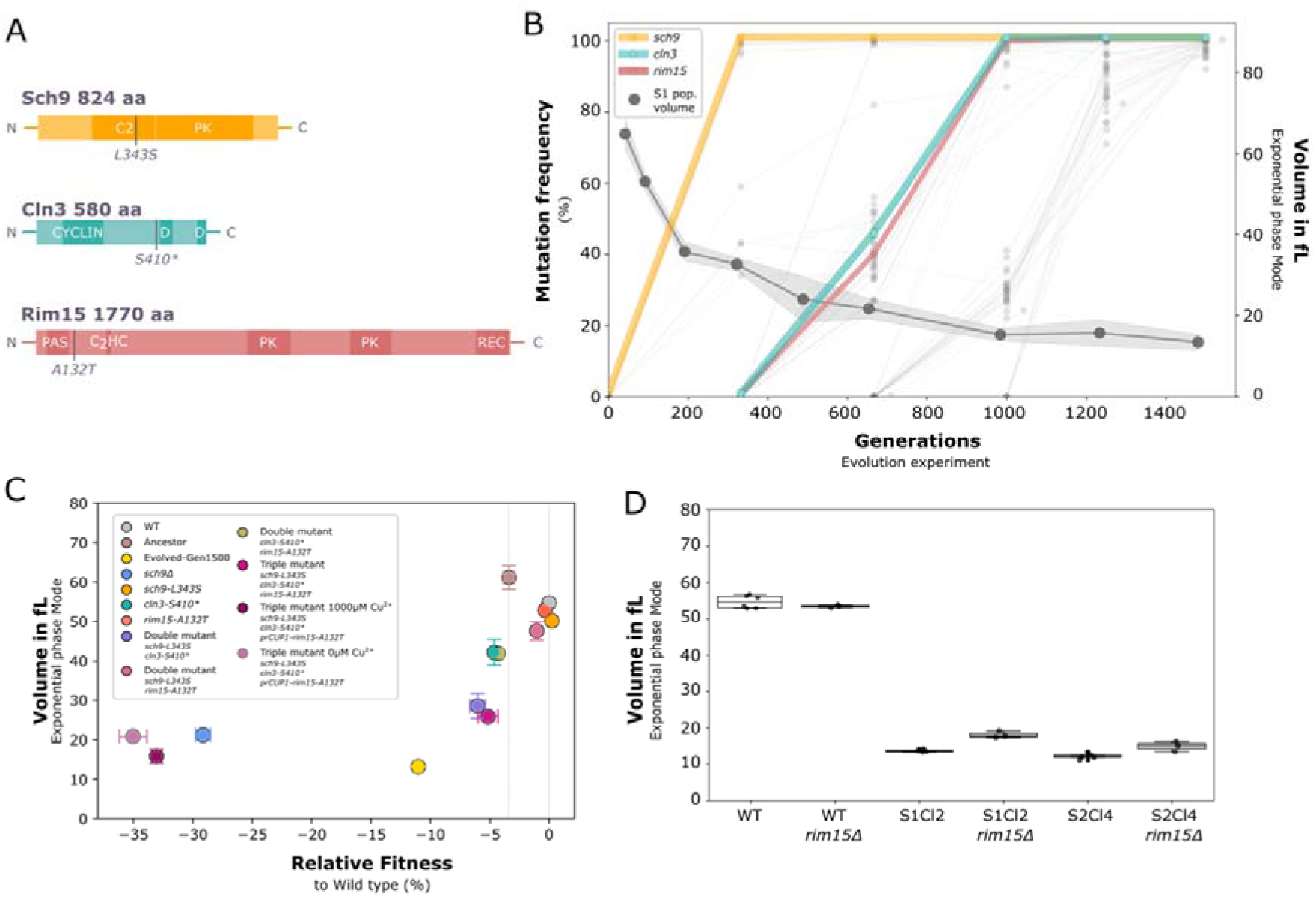
Genetic basis of the cell size phenotype. (A) Schematic of cell size–regulating genes. Protein domains are indicated, and mutations identified during evolution are marked: PK, kinase catalytic domain; D, consensus disorder prediction; PAS, N-terminal PAS domain; C2HC, CCHC-type zinc-finger domains; REC, C-terminal receiver domain. (B) Allele-frequency trajectories of size-associated mutations. Only variants reaching ≥80% frequency by generation 1,500 are shown; causative mutations are highlighted. The right axis shows the S1 population volume (circles) over time. (C) Relationship between relative fitness and modal cell volume. Each point represents a strain (wild type, ancestor, evolved populations, or engineered mutants). Fitness (x-axis) and cell volume (y-axis) are shown; error bars indicate SD. (D) Modal cell volume following deletion of *RIM15* in the wild-type background and in final evolved clones from exponential-phase cultures. Box plots show medians, interquartile ranges, whiskers (1.5× IQR).

To test the causality of these mutations, we reconstructed the *cim1*, *cln3*, and *sch9* alleles in a WT background. Deletion of *CIM1*, used to mimic the observed frameshift mutations, did not alter cell size (Figure S6A), suggesting a potential adaptive role in other cellular features or fitness. In contrast, mutations in *CLN3* and *SCH9* individually caused a modest but significant decrease in cell volume (Figure 6C). Combining the two mutations produced synergistic effects, with the double mutant exhibiting a three-fold reduction in modal volume (28.5 fL; Figure 6C).

Truncations of the C-terminal region of Cln3 confer a gain-of-function phenotype by increasing protein stability and cyclin abundance (65). Loss-of-function mutations in *SCH9* have likewise been associated with reduced cell size, typically attributed to its role in ribosome biogenesis (48). However, the elevated ribosome biogenesis observed in evolved lines (Figure 3G), together with their sustained fitness (Figure 3B–C), indicates that *sch9-L343S* is a separation-of-function allele that affects cell size through a mechanism distinct from ribosome biogenesis.

Recent work suggests that Sch9 can also regulate cell size via the Greatwall kinase Rim15, albeit under conditions of low CDK activity (67). This pathway is highly conserved among eukaryotes and controls cell cycle progression in metazoans through endosulfine-mediated inhibition of the phosphatase PP2A (68, 69). While it links nutrient sensing to cell size in *S. pombe* (70), in *S. cerevisiae* it has primarily been implicated in the regulation of quiescence (71). We identified mutations across multiple components of this pathway, including a fixed mutation in *RIM15*, as well as lower-frequency mutations in the promoter of the endosulfine *IGO1* and in *CDC55*, which encodes a regulatory subunit of PP2A (72). Reconstruction of the evolved *RIM15* allele (*rim15-A132T*) resulted in a modest, non-significant reduction in cell volume (Figure 6C). However, the distribution of mutations across the pathway (*sch9-L343S*, *rim15-A132T*, *igo1-T-75G*, and *cdc55-L159P*; Figure S6B) suggests that the Greatwall pathway activity was modulated during evolution. To test this hypothesis, we engineered a copper-inducible tunable promoter (*prCUP1*) to control expression of *RIM15* and *rim15-A132T*. Copper titration revealed a clear dosage-dependent effect: basal expression reduced cell size, and overexpression induced further shrinkage in WT backgrounds (Figure S6B). Notably, overexpression of *rim15-A132T* in the *sch9-L343S cln3-S410\** background further reduced modal cell volume, matching that of the evolved lines (16 ± 0.1 fL, mean ± SD; Figures 6C and S6C–F). Consistent with increased Greatwall pathway activity, deletion of *RIM15* increased exponential-phase cell volume in evolved clones but had little effect in WT (Figure 6D and Figure S7A). In stationary phase, however, *RIM15* deletion increased cell size in both backgrounds, with a stronger effect in evolved cells (Figure S7B-D). Furthermore, combined deletion of *RIM15* and *cln3*Δ showed synergistic effects on increasing WT cell size in stationary phase (Figure S7E-J).

Importantly, despite their reduced size, all reconstructed strains carrying evolved alleles retained near-WT fitness (Figure 6C). Although enforced expression of *rim15-A132T* phenocopied the size of evolved clones in the *sch9-L343S cln3-S410\** background, it imposed a fitness penalty. This penalty is unlikely to arise from reduced cell size itself, as fitness improved while size decreased further at higher copper concentrations (Figure 6C), and instead points to deleterious consequences of constitutive Rim15 activation across the cell cycle. These observations suggest that, rather than simply increasing pathway output, evolution selected for combinations of mutations that tune Greatwall signaling in a context-dependent manner to minimize fitness trade-offs.

These results demonstrate that cell size in the evolved lines is dependent on Rim15 dosage and support a model in which, together with a hyperactive G1 cyclin Cln3, increased Greatwall pathway activity contributes to reduced cell volume, although the precise molecular mechanisms remain to be defined.

## DISCUSSION

How can evolution produce large shifts in cell size without incurring the detrimental effects associated with even small deviations from the norm? In development, abrupt changes often arise through ploidy increase or reduction (73). Linear scaling relationships between DNA content and cell volume are also observed across diverse taxa (74). Ploidy increases are important in plant evolution and likely account for part of the variation in cell size among related species (75). Yet, genome duplications alone cannot explain the wide diversity of cell sizes, as they also create barriers to sexual reproduction and would leave more pervasive genomic signatures if they were the main driver. Cells must therefore be able to modify size independently of DNA content, at least initially, especially under gradual selection for divergent sizes.

Decades of studies on eukaryotic cell cycle regulation have identified the main principles allowing cells to regulate their size, as well as many molecular players involved (reviewed in (11, 76). Mutations in many such players have been demonstrated to produce altered sizes (35, 36, 77). It is therefore intuitive to imagine that evolution could modulate cell size by fine-tuning these molecular players. Yet most such mutations reduce fitness, and it remains unclear which, if any, combinations of the dozens of known regulators could provide viable evolutionary paths to new cell sizes.

We addressed this question experimentally by imposing selection on *S. cerevisiae* cell size and fitness. Over 1,500 generations, cells shrank four-fold in modal volume compared to WT, without changes in ploidy and with minimal fitness costs. We show that miniaturized cells maintain an efficient and evolutionarily stable cell size homeostasis, despite a significantly altered cell cycle progression characterized by the near absence of a G1 phase. Whole genome sequencing revealed a complex genomic landscape, due to the mutator allele employed, and likely linked to the many direct and indirect changes cells experienced during the miniaturization. Nevertheless, re-engineering putative adaptive alleles showed that simultaneous perturbations of the G1 cyclin Cln3 and the TOR effector Sch9 recapitulate much of the evolved phenotype with minimal fitness cost (Figure 6C). Although consistent with a gain-of-function effect from C-terminal truncation of Cln3, our data do not support a primary role for Sch9 in size control via ribosome biogenesis (Figure 3G). We argue instead for its involvement as an inhibitor of Rim15. Together, manipulations in the G1 cyclin and Greatwall kinase signaling cascades produced a six-fold range within an otherwise constant genome, capturing up to 70% (221 / 315) of the natural variation in size seen across the *Saccharomycotina* over 400 million years of evolution (Figure 6C, S7E-J and Figure 7A).

**Figure 7.**
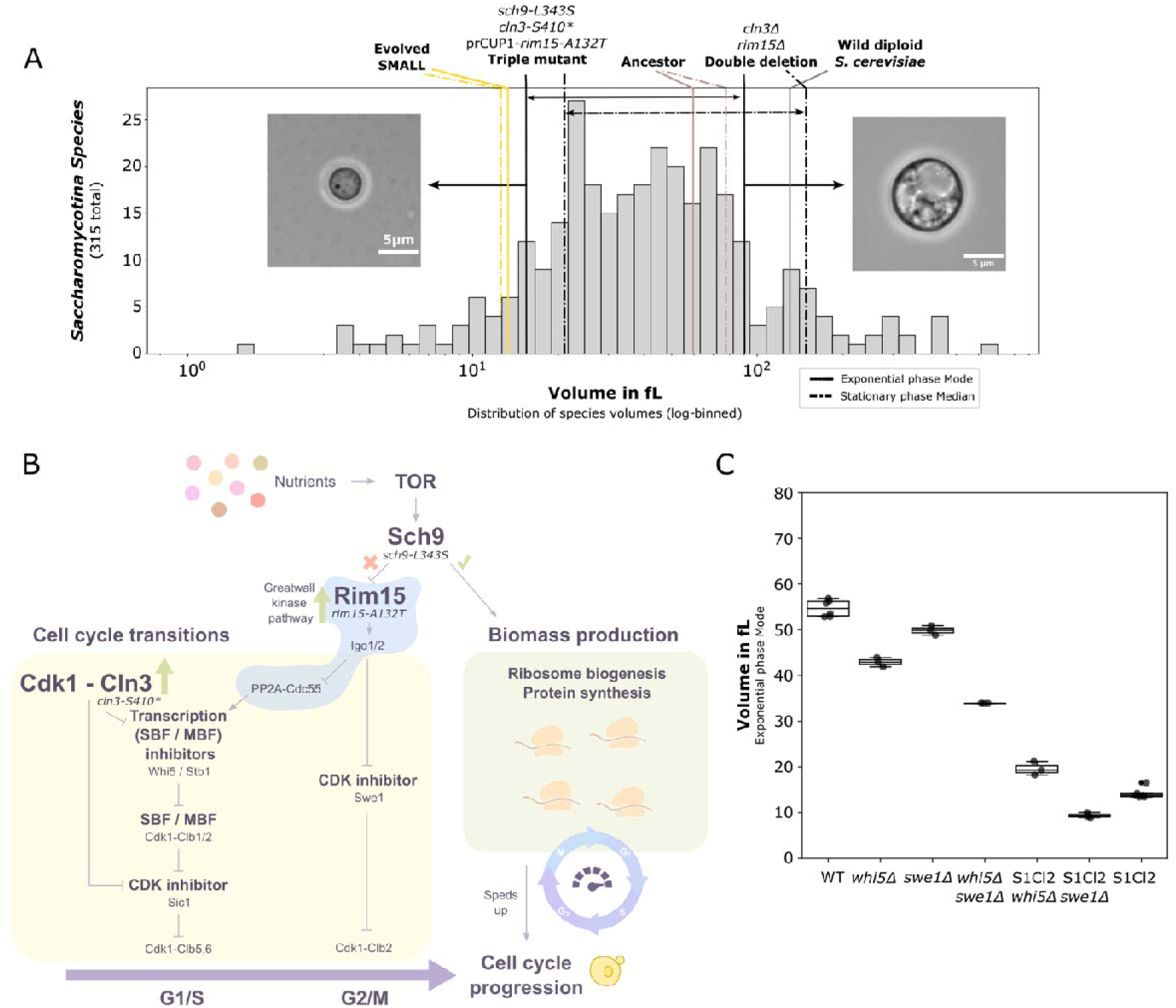
Speculative model of the effect of the mutations identified on cell cycle regulation. (A) Distribution of cell volumes across the *Saccharomycotina* subphylum (derived from (83)). Highlighted are the triple mutant (*sch9-L343S cln3-S410*\* *prCUP1-rim15-A132T*) under a copper-inducible promoter and the *CLN3* and *RIM15* double deletions, showing shifts in size relative to natural variation. (B) Modal cell volume of deletion of *WHI5* and *SWE1* in the wild-type background and in final evolved clones from exponential-phase cultures. Box plots show medians, interquartile ranges, and whiskers (1.5× IQR). (C) In wild-type (WT) cells growing under nutrient-rich conditions, TOR is active, promoting ribosome biogenesis and biomass accumulation (right, green rectangle) while delaying cell cycle transitions (left, orange rectangle). This results in fast-growing, large cells. Under nutrient limitation or in TOR-related mutants, Sch9 repression of Rim15 is relieved, leading to earlier cell cycle transitions. However, the accompanying reduction in biomass production limits growth and fitness. We propose that the combined effect of the identified mutations accelerates cell cycle transitions through truncations in Cln3, loss of Sch9 inhibition, and gain-of-function mutations in the Greatwall kinase cascade, while maintaining ribosome biogenesis and biomass production. As a result, cells grow rapidly but, by spending less time in interphase, remain smaller than their WT counterparts.

Three questions arise. First, how can our evolved cells reach sizes smaller than those achieved previously through gene deletions? We propose that evolution selected alleles with modest individual effects, but that act synergistically on cell size by working on multiple mechanisms simultaneously. We speculate the following model to account for the phenotype observed (Figure 7B): Truncations in *CLN3* stabilize the protein, increasing *CDK1–Cln3* activity and phosphorylation of the transcriptional inhibitor Whi5 (78). In parallel, loss of Sch9 inhibition and gain-of-function mutations in the Greatwall kinase cascade inhibit the phosphatase PP2A–Cdc55, further driving Whi5 phosphorylation and nuclear exclusion (67). But why would this lead to a volume smaller than *whi5Δ*, where the inhibitor is absent? First, CDK1-Cln3 levels could simultaneously hyper-phosphorylate the transcriptional regulator Stb1 (20, 79, 80) and accelerate the degradation of the CDK1 inhibitor Sic1 (81–83), eliminating the G1 phase. Furthermore, *RIM15* activity in mitosis has been reported to destabilize the CDK1 inhibitor and size/morphogenesis checkpoint Swe1 (84), offering an additional route to reduced volume. To test this model, we deleted *WHI5* and *SWE1*, inhibitors of the G1/S and G2/M transitions, respectively, in evolved clones (Figure 7C and Figure S8A). *WHI5* deletion increased cell size in evolved clones (19.3fL), whereas *SWE1* deletion caused a stronger size reduction than in WT (9% vs 33% decrease), producing cells with a modal volume of ∼9 fL. These results indicate that *whi5Δ* no longer contributes to size decrease in evolved lines, while Swe1 plays a more prominent role, suggestive of cell size regulation occurring in G2/M rather than G1/S.

The second question regards what allows these miniaturized cells to retain high fitness. Eukaryotic cells physiologically shrink upon nutrient scarcity (49). Under these conditions, cells may find it advantageous to increase population-wide survival by dividing at a smaller volume while making a parsimonious use of the resources left by reducing ribosome biogenesis. The deletion of *SCH9* hard-wires this response, resulting in small cells with severe fitness defects due to strongly decreased protein production. Here, we show that our evolved lines do not shrink further in poor-quality nutrients. Because we achieved even smaller cells through the deletion of Swe1, we conclude that the evolved lines have genetically altered the mechanism linking nutrient quality to cell size, rather than reaching a minimal size limit. Although such a mechanism has been proposed to operate through the modulation of ribosome biogenesis, our results show that ribosome biogenesis remains efficient in our evolved lines. This indicates that the effect of nutrients on cell size has become uncoupled from their effect on biomass production. This decoupling explains the high fitness of these evolved miniaturized cells and challenges current models of nutrient control of cell size.

Finally, have the evolved cells reached the lower size limit for *S. cerevisiae*? Gain-of-function mutants of the G1 cyclin Cln3, when grown in glycerol, approach the size of our evolved lines (16 fL, (53)). This observation supports our model in which the evolved lines combine size effects from the G1 cyclin pathway with those mediated by nutrient signaling. However, we argue that this combined effect does not represent the lower physical limit, as additional deletion of *SWE1* further reduces the modal cell volume to ∼9 fL. The long-term continuation of this evolutionary experiment will help determine whether, and how, a physical lower size limit can be reached experimentally.

In summary, our work shows that large deviations in cell volume can be achieved while maintaining cellular fitness and size homeostasis. We show that this phenomenon is accompanied by changes in cell cycle regulation and in the nutrient-dependent control of cell size. To achieve this phenotype, we propose a mechanism based on the simultaneous gain of function of the G1 cyclin and Greatwall kinase signaling pathways. In this model, increased Greatwall pathway activity arises, at least in part, from the relief of Sch9-mediated inhibition, thereby linking nutrient signaling to cell cycle control. We speculate that this model identifies a simple yet powerful route for evolution to achieve large deviations in volume without altering ploidy. Future studies should test the generality of the principles we identified and determine how broadly it underpins size diversity across the tree of life. The range of cell sizes generated within a single species, while maintaining high fitness and stable ploidy, also offers a framework to study subcellular scaling and the limits of cell size.

## MATERIAL AND METHODS

All strains used in this study were W303 derivatives. Yeast cultures were propagated using standard techniques (85). Unless otherwise stated, experiments were performed in standard rich medium (YPD). Experimental evolution followed the procedure described in (86), with the addition of the size-selection step by sorting detailed in the SI Appendix. Growth-curve measurements and fitness assays were performed as described in (87). Samples for cell-cycle analyses were processed as in (87) and analyzed as in (86). Whole-genome sequencing and analysis followed procedures described in (88). A full description of all methods is provided in the SI Appendix, Methods.

### Data Availability

The genomic datasets used within this publication were deposited at European Nucleotide Archive (PRJEB101156). Scripts used for data analysis are available at the GitHub repository (https://github.com/FumaLab/Garona2025). The raw data corresponding to each figure are provided in the Supplementary file Raw Data–Figures. Expanded view data, supplementary information, and appendices are available for this paper. Raw data from Coulter Counter measurements is available at Zenodo (DOI: 10.5281/zenodo.19254220).

### Authors contribution

M.F. conceived the project. A.Ga. and M.F. designed the research, performed the experiments, and analyzed the data. A.Gi. contributed data analysis for Figure 4B–C. M.V.L. contributed experimental work for Figure 5C, E. A.Ga. and M.F. wrote the original manuscript draft. All authors edited and commented on the final version of the manuscript.

## Supporting information

RawData-Figures

## Acknowledgments

We thank Mina Abrantes, Joana Boas, Filipa Cantanhede, Olga Carreira, João Martins, and Ana Quendera for experimental assistance; members of the Genome Maintenance and Evolution lab for support and discussions; and Gabriel Neurohr, Andrew W. Murray, Paula Gonçalves, Partha Pratim Chakraborty, and Mariana Natalino for manuscript review. We acknowledge the Flow Cytometry, Electron Microscopy, Bioimaging, and Genomics Facility of the Gulbenkian Institute for Molecular Medicine for technical support, and Beatriz Tomaz, Erin Tranfield, and Muriel Christel Mari for assistance with establishing electron microscopy sample preparation. We also thank Giorgio Tallarico and Marco Cosentino Lagomarsino for advice on quantifying the ribosomal fraction of the proteome. This work was supported by the Gulbenkian Foundation and the Gulbenkian Institute for Molecular Medicine (A.Ga., M.V.L., M.F.); Human Frontier Science Program (HFSP, RGEC28/2023; M.F., A.Gi.); European Molecular Biology Organization (EMBO, IG 5349-2023; M.F.); Fundação para a Ciência e a Tecnologia (FCT, 2023.09068.CEECIND; M.F.); and the National Institute of General Medical Sciences (NIGMS), National Institutes of Health (NIH), USA (1R35GM147493; A.Gi.).

## Notes

### Competing Interest Statement

The authors have declared no competing interest.

### Summary of Updates

Revised manuscript to include measurements performed in exponential phase for all strains; figures updated to incorporate exponential-phase data; supplemental files revised.

