## Appendix for "Experimental evolution of cellular miniaturization reveals a putative mechanism for cell size evolution"

**SI APPENDIX**

### METHODS

#### Strains and culture conditions

All strains used in this study were derived from *Saccharomyces cerevisiae* W303 (*leu2-3,112 trp1-1 can1-100 ura3-1 ade2-1 his3-11,15, RAD5*$^{+}$) and are listed in Table S1. The haploid strain yMF1001 (*ADE2; leu2-3,112; ura3-1; trp1-1; RAD5*$^{+}$*; yel068c::CAN1/URA3; prACT-Citrin-LEU2; pol3-L523D*) was used as the ancestral strain in the evolution experiments. This strain carries the *pol3-L523D* mutator allele, which lacks the proofreading activity of the main replicative polymerase and increases the mutation rate by approximately 100-fold (1). Yeast propagation was carried out using standard techniques (2). Unless otherwise specified, all experiments were performed in standard rich medium (YPD), consisting of 1% yeast extract, 2% peptone, and 2% D-glucose, supplemented with 0.01% adenine and tryptophan. For the evolution experiments, the medium was further supplemented with penicillin and streptomycin. Auxotrophic selection and complementation assays were performed in Synthetic Complete (SC) medium or in the appropriate dropout variants. SC medium consisted of 10% (v/v) 10× Yeast Nitrogen Base (YNB) without amino acids, supplemented with 0.768 mg/L each of arginine, isoleucine, lysine, phenylalanine, and tyrosine; 1.152 mg/L aspartic acid; 0.256 mg/L methionine; 1.536 mg/L threonine; 2.176 mg/L valine; and 2% D-glucose. Depending on the auxotrophic markers required, SC medium was further supplemented with 0.025 mg/mL each of histidine, uracil, tryptophan, and adenine, and with 0.05 mg/mL leucine. Either *Saccharomyces cerevisiae* or *Escherichia coli* DH5α were used for strain construction. Yeast strains were generated using the standard lithium acetate (LiAc) transformation method. *E. coli* DH5α was routinely cultured in lysogeny broth (LB) medium at 37 °C with shaking at 250 rpm or on LB agar plates. DNA concentrations were measured using either a NanoDrop spectrophotometer (Thermo Fisher Scientific) or a Qubit fluorometer (Thermo Fisher Scientific). All construct assembly steps were verified by PCR, restriction enzyme digestion, or Sanger sequencing. The proofreading Phusion DNA polymerase (Thermo Fisher Scientific) was used for all PCR-based cloning.

**Arrest and release of yeast cell cycle**

Logarithmically growing cells were arrested in G1 phase by incubation in YPD medium supplemented with α-factor (3 µg/mL) for 2 h. Cell cycle arrest was confirmed by examining cell morphology. Following arrest, cells were washed three times with room-temperature YP medium (lacking glucose) and resuspended in fresh YPD medium to allow cell cycle re-entry. Cultures were incubated for 4h to monitor recovery of normal cell cycle progression. Samples were collected periodically to assess cell morphology, size, and DNA content.

**Fluorescence-activated cell sorting (FACS)**

Cell sorting was performed using a FACSAria cytometer (BD Biosciences) with PBS as the sheath fluid. A 70-µm nozzle was used, and the sorting mode was set to “Purity.” The flow cytometer voltage for SSC and FITC channels were set at 311 and 453 respectively. Forward scatter (FSC) voltages were adjusted between 220 V and 400 V to optimize resolution across cell sizes. The flow rate was set to 1, and the cell concentration was adjusted to maintain an event rate below 9000 events $s^{-1}$. Cultures were diluted as needed in 50 mM Tris-HCl (pH 7.5) and sonicated for 50 s at 14% amplitude using a Branson SLPe Digital Sonifier. Gating was performed on bivariate dot plots to identify single yeast cells. The initial gate was set on FITC-A versus FSC-H to select events with high fluorescence in the FITC channel, corresponding to expression of the fluorescent reporter. Cells were then gated on SSC-A versus FSC-A. Doublets were excluded in two steps to prevent sorting of aggregated cells using FSC-W versus FSC-A and FSC-H versus FSC-A gating. Cells were finally sorted based on FSC-A, used as a proxy for cell size. The smallest 7% of the population (corresponding to small-sized cells) was defined as the SMALL population.

Each day, 100,000 cells from the lower 7% were collected. Sorted cells were recovered in Eppendorf tubes containing 500 µL of YPD medium and subsequently transferred to glass tubes containing 10 mL of fresh YPD.

**Clone selection by Forward Scatter (FSC-A)**

Single colonies from each evolved population at 1,500 generations were picked from YPD plates and inoculated into liquid YPD medium, followed by overnight incubation at 30 °C. Cultures were then diluted and analyzed using an LSRFortessa flow cytometer (BD Biosciences). Forward scatter area (FSC-A) was used as a proxy for cell size, while doublets and debris were excluded based on FSC-H versus FSC-A and SSC-A gating. For each clone, at least 10,000 singlet events were collected at a low flow rate. The median FSC-A value per clone was used for comparative analyses. Clones S1Cl2 and S2Cl4 were selected to proceed with the experiments requiring a homogeneous genetic background.

**Evolution Experiments**

The size selection experiment was initiated using the haploid ancestral strain yMF1001, grown overnight in rich YPD medium. At the start of the experiment, the ancestral culture was sorted by FACS to generate three distinct populations (S1-3 lineages). Cells were selected based on their small forward scatter area (FSC-A) values relative to the rest of the population, as described previously.

Populations were cultivated in glass tubes on roller drums at 30 °C and grown for 20 h. Each day, cultures were sorted, and 100,000 sorted cells were resuspended in 10 mL of fresh YPD medium, allowing approximately 11–13 generations per cycle. Populations were propagated for a total of 116 cycles (≈1,500 generations). At each cycle, 800 µL of each evolving population was mixed with 800 µL of 30% (v/v) glycerol and stored at –80 °C for long-term preservation. The evolution experiment in the absence of size selection was performed using the wild-type (WT) strain, the ancestral strain, and two single clones isolated at the end of the size-selection evolution experiment. In addition, a *sch9Δ* deletion mutant was included. Each strain was inoculated into glass tubes containing 10 mL of YPD medium. After 24 h of growth, three independent replicates were established per strain, yielding a total of 12 evolving populations. Cultures were incubated under the same conditions as in the original evolution experiment, at 30 °C with continuous rotation, and grown for 24 h each cycle. Daily passages were performed by transferring 10 µL of the previous culture into 10 mL of fresh YPD (1:1000 dilution), allowing approximately 10 generations per cycle. Populations were propagated for 15 cycles (≈150 generations). Every five cycles (≈50 generations), 800 µL of each population was mixed with 800 µL of 30% (v/v) glycerol and stored at –80 °C for long-term preservation.

**Whole genome sequencing**

Genomic DNA was extracted by standard ethanol precipitation from 1 mL of yeast culture. Population samples collected at generations 333, 666, 1000, 1250, and 1500 were sequenced to identify genomic variants. Sequencing was also performed on populations derived from bulk segregant analysis.

Sequencing depths were approximately 25x for individual clones, 200x for population samples, and 500x for bulk segregant populations. Genomic DNA libraries were prepared following the protocol described by Koschwanez *et al.* (3) using the Nextera DNA Library Prep Kit (Illumina, RRID:SCR_010233; San Diego, CA, USA). DNA library concentration and quality were assessed with a GloMax® Plate Reader (Promega) and the Quant-iT™ PicoGreen® dsDNA Assay (Thermo Fisher Scientific). Paired-end sequencing was performed on an Illumina NextSeq 2000 platform.

**Convergent evolution on genes**

This method was performed based on the analysis of Fumasoni and Murray (4). Briefly, it relies on the assumption that those genes that have been mutated significantly more than expected by chance alone, represent cases of convergent evolution among independent lines. The mutations affecting those genes are therefore considered putatively adaptive. Per-base mutation rates were calculated as the number of mutations in coding regions divided by the total coding sequence length of the yeast genome in bp, for a given genetic background (including 1000 bp per ORF to account for regulatory regions).

$$\lambda = \frac{SNPs + indels}{\text{Coding Base Pairs}}$$

If the mutations were distributed randomly in the genome at a rate λ, the probability of finding n mutations in a given gene of length N is given by the Poisson distribution:

$$P(n\ mutations | gene\ of\ length\ N) = \frac{(\lambda N)^n e^{-\lambda N}}{n!}$$

For each gene of length N, we then calculated the probability of finding n mutations if these were occurring randomly:

$$P(\geq n\ mutations | gene\ of\ length\ N) = \sum_{k=n}^{\infty} \frac{(\lambda N)^n e^{-\lambda N}}{n!} = 1 - \frac{\Gamma(n, \lambda N)}{(n-1)!}$$

(where Γ is the upper incomplete gamma function) which gives us the P value for the comparison of the observed mutations with the null, Poisson model. To decrease the number of false positives, we then performed multiple-comparison corrections. The Bonferroni correction was used. The custom pipeline used for the data analysis is available on GitHub.

**Bulk segregant analysis**

Bulk segregant analysis (BSA) is an experimental approach used to identify genetic loci associated with a specific phenotype (3, 5). In this study, BSA was performed by crossing a clone from the evolved population exhibiting the phenotype of interest with a WT strain lacking that phenotype. The resulting diploids were sporulated, allowing random segregation of the mutations present in the evolved clone. Haploid segregants were selected for both growth and the phenotype of interest—in this case, small cell size—over approximately 40–50 generations by sorting as described for the evolution experiment set-up. The fittest cells carrying the causative mutations became enriched in the population during selection. Segregants displaying the target phenotype were subsequently pooled, and their genomic DNA was extracted, sequenced, and analyzed to identify mutations associated with the phenotype. BSA was performed as follows. One clone (*MATα ADE2; leu2-3,112; ura3-1; trp1-1;* $RAD5^{+}$*; yel068c::CAN1/URA3; prACT-Citrin-LEU2; pol3-L523D*) from each evolved population was selected as described above (see *Clone selection by forward scatter*) and mated with a haploid MATa WT strain (*yMF0473; W303 background: ade2-1; can1-100; leu2-3,112; his3-11,15; ura3-1; trp1-1;* $RAD5^{+}$*; pSTE2-URA3::KanMX4*). In this WT strain, the endogenous *URA3* promoter was replaced with the *STE2* promoter, which is active only in MATa cells, allowing selection for MATa spores after meiosis since neither MATα haploids nor MATa/MATα diploids can express *URA3* under the *STE2* promoter. Mating was carried out on YPD plates by mixing cells from both strains. The crosses were incubated overnight at 30 °C, after which cells were scraped from the plates and grown in double-selective synthetic medium lacking histidine and tryptophan (–HIS –TRP) to select for diploids. Diploid cultures (100 mL) were grown to saturation, centrifuged, and resuspended in synthetic medium containing 2% potassium acetate (KAc) to induce sporulation. Cultures were incubated overnight (12–15 h) at 30 °C, pelleted, and resuspended in water containing 2% KAc. Sporulation was continued for 7 days at room temperature (23–25 °C) with agitation on a roller drum. Sporulation efficiency was verified microscopically and was approximately 15%, yielding 0.5–1 × $10^9$ spores. To digest asci, 100 mL of the sporulated culture was pelleted and resuspended in 5 mL of water containing 2500 units of Zymolyase (Zymo Research). The suspension was incubated with shaking at 30 °C for 1 h, with intermittent vortexing. When most tetrads were separated, 40 mL of water and 5 mL of 10% Triton X-100 were added, followed by 1 min of sonication to disperse remaining aggregates. Spores were collected by centrifugation (6000 rpm, 1 min) and resuspended in haploid selective medium lacking uracil (–URA). After 48 h of growth, spores were subjected to fluorescence-activated cell sorting (FACS) to select for cells with small size and high fitness, as described previously. This selection regimen, analogous to that used during the experimental evolution, enriches for cells displaying the target phenotype. After three rounds of selection, genomic DNA was extracted from the final saturated cultures and used for library preparation and high-coverage whole-genome sequencing, as described above. A custom pipeline was used to identify alleles segregating with the evolved phenotype. Variants present at frequencies greater than 50% were considered non-randomly segregating, consistent with inheritance from the evolved parent in the diploid cross (haploid evolved × haploid WT). A second BSA was performed with clones isolated from the first BSA, to narrow down the set of candidate mutations: the first bulk yielded approximately 30–80 segregating mutations, whereas the second bulk reduced this number to around 10 candidate mutations.

**Fitness assays**

The relative fitness ($\omega$) of WT, ancestral, evolved, or constructed strains relative to a WT reference strain was determined through direct pairwise competition experiments, each performed in triplicate. The reference strain was haploid *S. cerevisiae* W303 (yMF1176: *can1; ADE2; leu2-3,112; his3-11,15; ura3-1; trp1-1; RAD5$^+$; yel068c::CAN1/URA3; prACT-Citrin-LEU2; prACT-Cerulean-KANMX*). All strains were preconditioned by growth to saturation in liquid YPD medium prior to competition. Competing strains were mixed at defined initial ratios (typically 1:1 or 20:80 depending on competing strain growth) and incubated at 30 °C for 24 h. Cultures were serially transferred through 1:1000 dilutions for three consecutive 24 h cycles, and the relative abundances of competing strains were quantified at each transfer. Population sizes were determined based on the number of fluorescent and non-fluorescent events detected by flow cytometry (LSRFortessa, BD Biosciences). The reference strain expressed a Cerulean fluorescent protein, which was absent in the WT, ancestral, evolved, and constructed strains, allowing strain discrimination. Relative fitness was estimated from changes in the ratio of competing strains over time. Specifically, the slope of a linear regression of the natural logarithm of the population ratio (fluorescent/non-fluorescent events) against the number of generations was used to calculate $\omega$. Relative fitness values were normalized to those of the WT or ancestral strains, depending on the comparison. Reported *P*-values were derived from two-tailed *t*-tests. The custom data analysis pipeline is available on GitHub.

**Cell cycle profiles**

Cell cycle analysis was performed on cells collected from exponentially growing cultures. Cells were harvested by centrifugation (2 min at 6000 rpm) and resuspended in 70% ethanol containing 250 mM Tris-HCl (pH 7.6). The suspension was sonicated for 10 s at 10% amplitude using a Branson SLPe Digital Sonifier and incubated for 1 h at room temperature. Cells were then washed with 50 mM Tris-HCl (pH 7.5), resuspended in the same buffer supplemented with 0.04 mg/mL RNase A (Sigma-Aldrich), and incubated at 37 °C for at least 1 h. Following RNase treatment, cells were pelleted and resuspended in 50 mM Tris-HCl (pH 7.5) containing 0.2 mg/mL proteinase K (GRiSP), and incubated for 30 min at 50 °C. After centrifugation, cells were resuspended in 50 mM Tris-HCl (pH 7.5). Samples were diluted 10–20-fold in 50 mM Tris-HCl (pH 7.5) containing 1 mM SYTOX Green and sonicated again for 10 s at 10% amplitude. Stained samples were analyzed on an LSRFortessa flow cytometer (BD Biosciences) using the FITC channel to quantify cellular DNA content. Flow cytometer settings were as follows: 10,000 events acquired per sample; FSC 340; SSC 200; FSC PMT 130; SSC(BV) 190; and FITC 535–590. Cell cycle profiles were analyzed and visualized using a custom Python script based on the *FlowCytometryTools* package (https://pypi.org/project/FlowCytometryTools/). The fraction of the doubling time spent by each sample in each cell cycle phase was calculated using RepliFlow (6).

**Growth curves**

Overnight cultures were diluted 1:300 (50 µL into 15 mL of total volume) into rich medium and grown at 30 °C in culture tubes with continuous shaking to minimize sedimentation. Cell density was measured every hour with a Coulter Counter (Multisizer 4e, Beckman Coulter). Time-series data were analyzed in Python v3.9 using a custom script based on *growth_curve_analysis.py* (https://github.com/nwespe/OD_growth_finder), and the “effective growth rate” mode was used to calculate growth rates.

**Mating type switch**

SMALL clones from evolved populations S1 and S3 were transformed with a *pGAL1-HO* (HO: Homothallic switching endonuclease) plasmid conferring nourseothricin resistance. Evolved clones from population S2 failed to switch mating type. Single transformant colonies were grown for 24 h at 30 °C in liquid YP medium supplemented with 2% raffinose to relieve repression of the *GAL1* promoter. The following day, cultures were diluted 1:10, and galactose was added to a final concentration of 2% to induce expression of HO endonuclease. After 90 min of induction to allow for mating-type switching, cultures were diluted 1:100 and plated onto YPD plates. Successful mating-type switches were verified by mating the resulting cells with tester strains of known mating type.

### Volumetric measurements

Cell volume was determined using a Coulter Counter (Multisizer 4e, Beckman Coulter), which measures particle size based on the Coulter principle—each cell passing through an electrolyte-filled aperture generates an impedance change proportional to its volume (7). Before measurement, cultures were briefly sonicated for 10–30 s at 10% amplitude using a Branson SLPe Digital Sonifier to reduce cell clumping. Cells were diluted in Coulter Counter cuvettes containing 10ml of Coulter ISOTON II solution. For exponential-phase measurements, cells were sampled 7 h after a 1:200 dilution, and 100 μL of sonicated culture was used. For stationary-phase measurements, cells were collected at 48 h after dilution, with 10 μL of sonicated culture taken for analysis. A 50 μm aperture tube was used. For all volume estimations, a minimum of 30,000 events were recorded, and particles smaller than 1.5 fL or larger than 2500 fL were excluded from analysis. Raw data are available at Zenodo (DOI: 10.5281/zenodo.19254220). Data analysis was performed using a custom Python pipeline available on GitHub.

### Quantification of modal volumes and of the width of cell volume distributions

Cell size measurements obtained with the Coulter Counter included noise contributions from debris and non-cellular particles. Such artifacts affect calculations of summary statistics such as the mean, median, mode and standard deviation. Therefore, we characterized each cell size distribution by two statistics robust to low-volume debris: the modal volume and the normalized mean excess above the mode.

The modal volume $v_{mode}$ was estimated as the location of the maximum of a kernel density estimate (KDE) of the cell volume distribution, restricted to volumes above a strain-specific debris cutoff (Fig. 4B). Because debris particles are smaller than intact cells, the mode of the cellular peak is unaffected by their presence as long as they are well-separated from the main distribution.

To quantify the width of the right tail of the size distribution, we computed the mean excess above the mode, defined as $M = E[v - v_{mode} | v \geq v_{mode}]$. This quantity measures the average distance of cells at or above the mode from the modal volume, i.e. the average "extra size" carried by cells in the right tail. Normalizing by the modal volume yields the dimensionless ratio $M/v_{mode}$ reported in Fig. 4B. This metric is a natural generalization of the right Half-Width at Half Maximum (HWHM) of the size distribution. The HWHM measures the distance from the mode to the point where the density falls to half its peak value: a single horizontal slice of the distribution. By contrast, it can be understood as the average of all such right half-widths across every density level $h$ from $0$ to $p(v_{mode})$: by the layer-cake identity, $\int_{v_{mode}}^{\infty} (v - v_{mode}) p(v) dv = \int_0^{p(v_{mode})} w(h) dh,$ where $w(h) = sup\{v \geq v_{mode} : p(v) \geq h\} - v_{mode}$ is the right semi-width of the distribution at height $h$ (Fig. 4C, right panel). Thus, $M$ captures the full shape of the right tail rather than its width at one arbitrary height, while remaining interpretable in the same units as the HWHM.

**RNA extraction**

Total RNA was extracted from yeast cultures with three biological replicates per strain. $1.5 \times 10^7$ cells were collected during exponential growth phase and RNA extracted with the YeaStar™ RNA Kit from ZymoResearch. RNA integrity was evaluated by the RQN parameter (RNA quality number) with the 5200 Fragment Analyzer (Agilent Technologies) according to the manufacturer's instructions. RQN was >6 in all the samples. RNA concentration was measured with the NanoDrop spectrophotometer (Thermo Fisher Scientific).

**Protein extraction**

Total protein was extracted from yeast cultures with three biological replicates per strain. $1 \times 10^8$ cells were collected at exponential growth phase and protein extracted with a Trichloroacetic Acid (TCA) based method. Cells were harvested by centrifugation for 3 min at 4,000 rpm. The supernatant was discarded, and cell pellets were resuspended in 2 mL of 20% TCA in 2 mL Eppendorf tubes. Cells were then centrifuged for 1 min at 13,000 rpm, and pellets were resuspended in 200 µL of 20% TCA. Glass beads were added to the suspension, leaving a small volume without beads, and samples were vortexed at 4°C for 3 min to lyse the cells. Subsequently, 400 µL of 5% TCA was added, and the mixture was centrifuged for 10 min at 3,000 rpm. The supernatant was carefully transferred to a fresh 1.5 mL Eppendorf tube. Pellets were allowed to dry completely on paper. Dried pellets were resuspended in 100 µL of bicinchoninic acid assay (BCA) compatible extraction buffer (50 mM Tris-HCl, pH ~7.5; 150 mM NaCl; 0.5% non-ionic detergent such as Triton X-100 or NP-40/IGEPAL) for downstream protein quantification. Protein concentration was determined using a BCA assay according to the manufacturer's instructions.

**Electron microscopy**

*Saccharomyces cerevisiae* cells were grown in YP medium supplemented with 0.5% D-glucose. This reduced glucose concentration was chosen according to yeast EM best practices to improve cell wall permeability. A total of 10–15 $OD_{600}$ units of cells were collected and centrifuged at 1000 g (≈3000 rpm) for 3–5 min at room temperature. The resulting pellet was resuspended and washed for 10 min with 10 mL of distilled water. This washing step is essential for autophagic body analysis, as it promotes the formation of a single large vacuole. Cells were then resuspended in 1.5 mL of freshly prepared ice-cold 1.5% $KMnO_4$ fixative and transferred into 1.5 mL microcentrifuge tubes. Tubes were filled with the fixative to eliminate air and incubated on a rotating wheel for 30 min at 4 °C. Cells were centrifuged at 1400 × g (≈4000 rpm) for 3 min at 4 °C, resuspended in 1.5 mL of ice-cold 1.5% $KMnO_4$, and incubated overnight on a rotating wheel at 4 °C. The following day, cells were washed five times with distilled water, centrifuging each time at 2200 × g (≈5000 rpm) for 3 min. Samples were then dehydrated through a graded acetone series (10%, 30%, 50%, 70%, 90%, 95%, and 100%), with each step lasting 20 min. Between each step, cells were centrifuged at 2200 × g (≈5000 rpm) for 4 min. After dehydration, samples were infiltrated with SPURR resin. The infiltration was performed sequentially as follows: (1) 1 part resin to 3 parts acetone for 1 h at room temperature; (2) 1:1 resin-to-acetone for 1 h; and (3) 3:1 resin-to-acetone for 1 h, with centrifugation (≈5000 rpm, 3 min) between each step. Finally, samples were incubated overnight in 100% resin. The following day, samples were centrifuged at 2200 × g (≈5000 rpm) for 5 min, the supernatant discarded, and a final infiltration in 100% resin was performed for 1 h at room temperature on a rotating wheel. Samples were transferred to conical embedding capsules, centrifuged (5000–7000 rpm, 3–5 min), the supernatant discarded, and the tubes filled with fresh 100% SPURR resin. Polymerization was carried out at 60–70 °C for at least 3 days. Polymerized samples were sectioned to a depth of approximately 10 μm using standard methods, and ultrathin sections (≈80 nm thick) were obtained with a diamond knife. Sections were imaged on a Tecnai G2 Spirit BioTWIN transmission electron microscope (FEI) equipped with an Olympus-SIS Veleta CCD camera. Three-dimensional image reconstruction was performed using Amira software (Thermo Fisher Scientific). Image stacks were aligned, and organelles were manually segmented and identified.

**Distribution of cell volumes across the *Saccharomycotina* subphylum**

Cell volumes were estimated from measured mean cell length and width (μm) from (8) by approximating each morphology with an idealized geometric solid. For each annotated shape class, a corresponding analytical formula (e.g., sphere, ellipsoid, cylinder, spherocylinder, cone, or frustum) was used to compute volume. When multiple shape annotations were present for a given cell type, volumes were calculated independently for each applicable geometry and averaged to obtain a single estimate. Invalid or non-physical measurements (e.g., negative or non-finite values) were excluded. All volumes were expressed in femtoliters, using the identity 1 $\mu m^3$ = 1 fL. The code used for these calculations is available in a public GitHub repository. (https://github.com/FumaLab/Garona2025).

**SUPPLEMENTARY FIGURES**

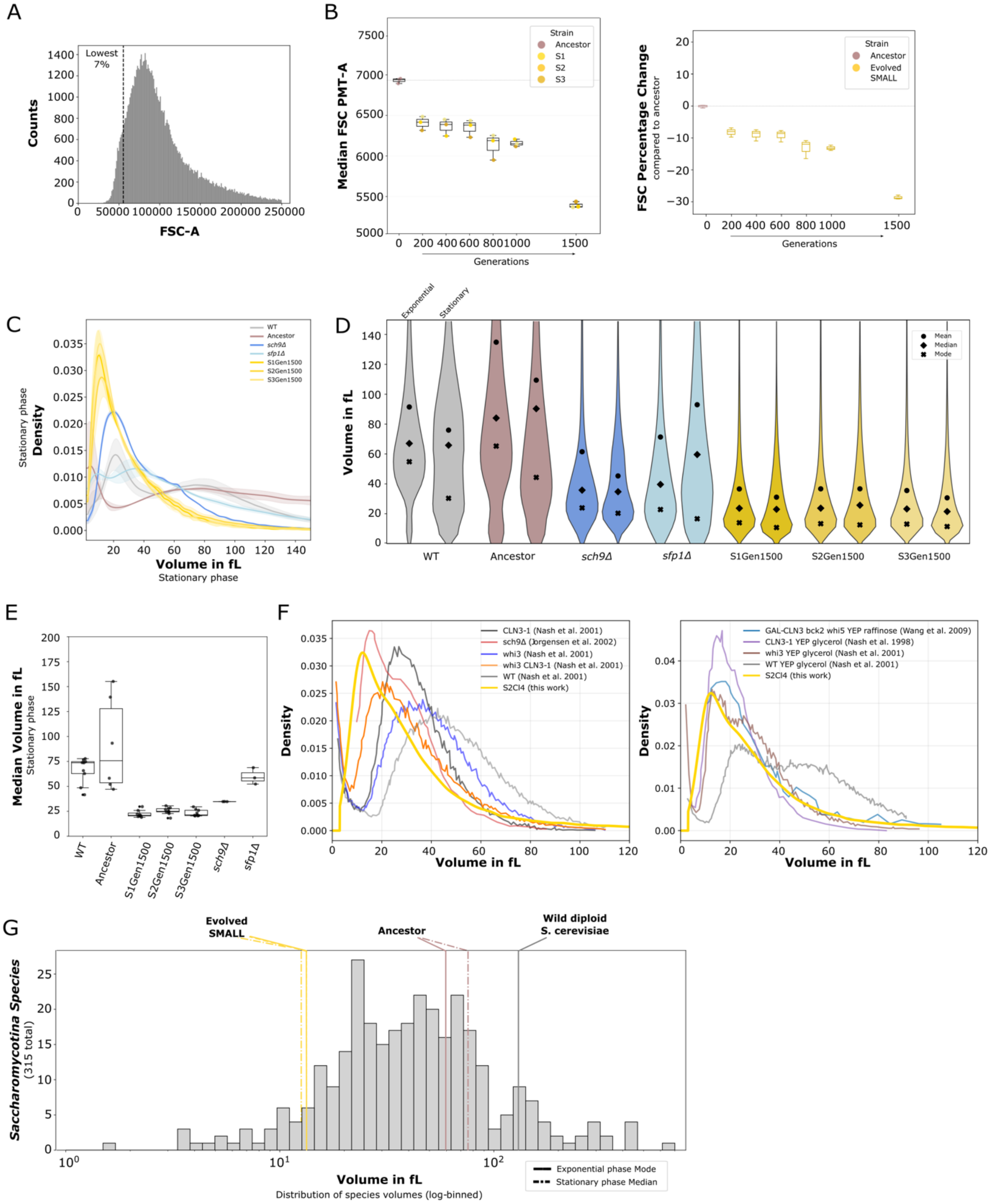


**Figure S1.** (A) Selection of the lowest 7% of the population based on forward scatter area (FSC-A). (B) Changes in median and relative FSC PMT-A during experimental evolution across populations. (C) Cell volume distributions in stationary-phase cultures. Smooth density curves represent kernel density estimates fitted to histograms of cell volumes measured by Coulter counter. For each strain, densities were computed per replicate and averaged to obtain a mean density curve. Shaded areas represent the standard deviation across replicates. (D) Cell size dynamics across growth phases. Violin plots show the average distribution of cell volumes (fL) measured in exponential and stationary phases. Summary statistics highlighted were derived from all individual measurements. (E) Median cell volume from measurements of stationary -phase cultures (Coulter counter). Box plots show medians, interquartile ranges, whiskers (1.5× IQR). (F) Cell volume distributions for the evolved clone S2Cl4 and previously reported small-size mutants measured under standardized conditions. Left: Cells grown in rich medium (YPD) and sampled during exponential phase. Right: Cells grown in nutrient-limiting media containing poor carbon sources (e.g., glycerol or raffinose) and measured during exponential phase. Cell size distributions for reference strains were digitized from published figures (9–12). (G) Distribution of cell volumes across the *Saccharomycotina* subphylum, with volumes estimated from cell diameters and shapes reported in (8). Only 29 of 315 species (9.21%) have a volume ≤14 fL, comparable to the evolved SMALL strain.

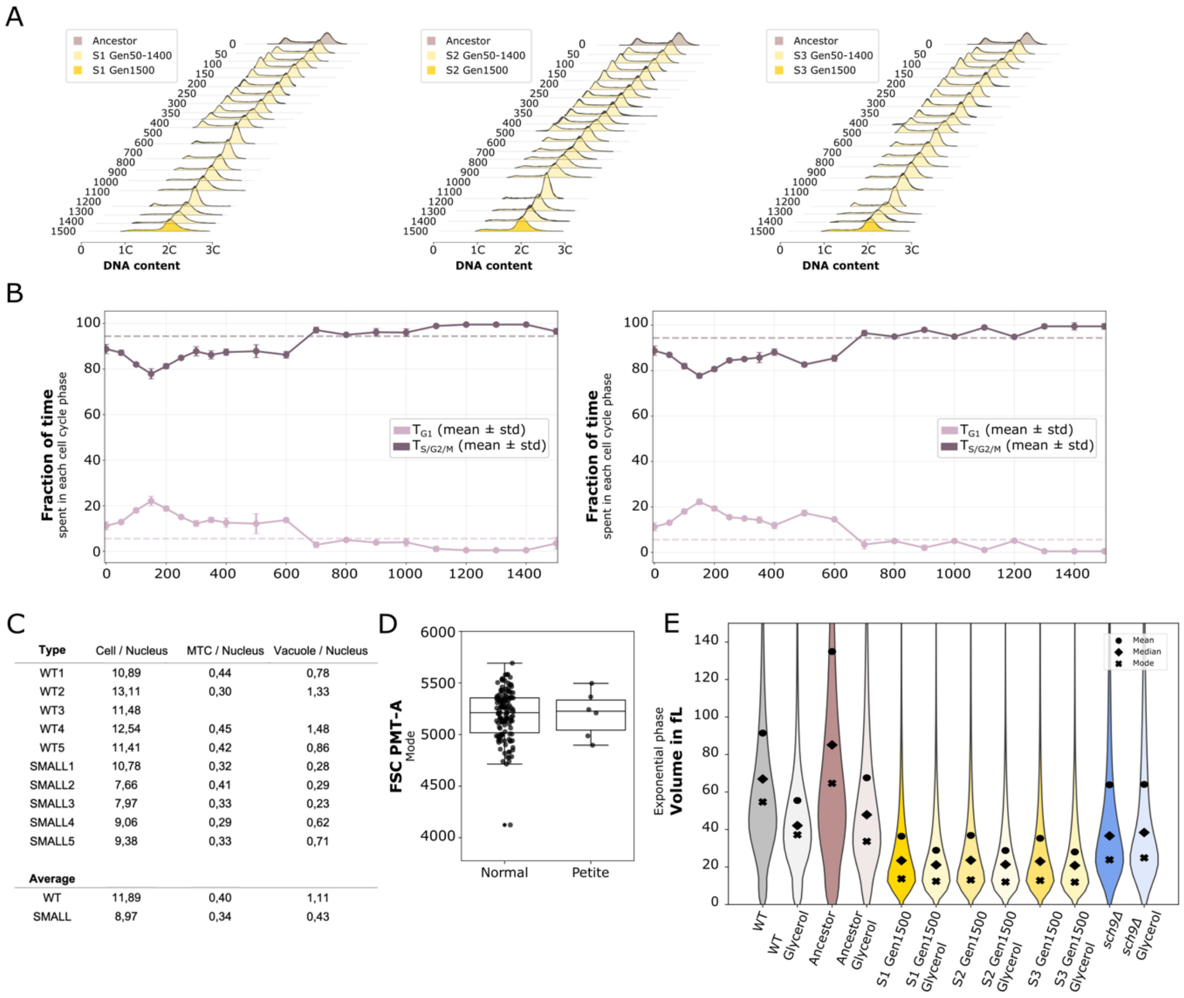


| Type | Cell / Nucleus | MTC / Nucleus | Vacuole / Nucleus |
|---|---|---|---|
| WT1 | 10,89 | 0,44 | 0,78 |
| WT2 | 13,11 | 0,30 | 1,33 |
| WT3 | 11,48 | | |
| WT4 | 12,54 | 0,45 | 1,48 |
| WT5 | 11,41 | 0,42 | 0,86 |
| SMALL1 | 10,78 | 0,32 | 0,28 |
| SMALL2 | 7,66 | 0,41 | 0,29 |
| SMALL3 | 7,97 | 0,33 | 0,23 |
| SMALL4 | 9,06 | 0,29 | 0,62 |
| SMALL5 | 9,38 | 0,33 | 0,71 |
| **Average** | | | |
| WT | 11,89 | 0,40 | 1,11 |
| SMALL | 8,97 | 0,34 | 0,43 |

**Figure S2.** (A) Cell cycle profiles of SMALL lineages S1–S3, shown without normalization to the G2 peak. DNA content was estimated using SYTOX Green staining (~10,000 cells per histogram). The x-axis shows arbitrary fluorescence units; the y-axis, relative cell frequencies. Solid lines denote mean profiles; shaded areas, SD (n = 3). Peaks for 1C (G1) and 2C (G2/M) DNA content are indicated. (B) Allocation of time to G1 and S/G2/M phases in S2 (left) and S3 (right) populations, inferred from DNA content profiles. Phase durations were calculated following (6). Bars show mean ± SD (n = 3). (C) Transmission electron microscopy (TEM) measurements of cell and organelle size, with nuclear size included as an internal control. The WT3 sample was excluded due to a sectioning artifact during sample preparation that compromised the nucleus area. (D) Mitochondrial functionality assessment. Among 137 isolated clones, 6 (4.3%) displayed mitochondrial defects. The y-axis shows FSC-PMT-A values indicating size similarity between petites and normal cells. (E) Cell size across carbon sources. Violin plots show the average distribution of cell volumes (fL) measured in rich media with glucose as carbon source or glycerol. Summary statistics highlighted were derived from all individual measurements.

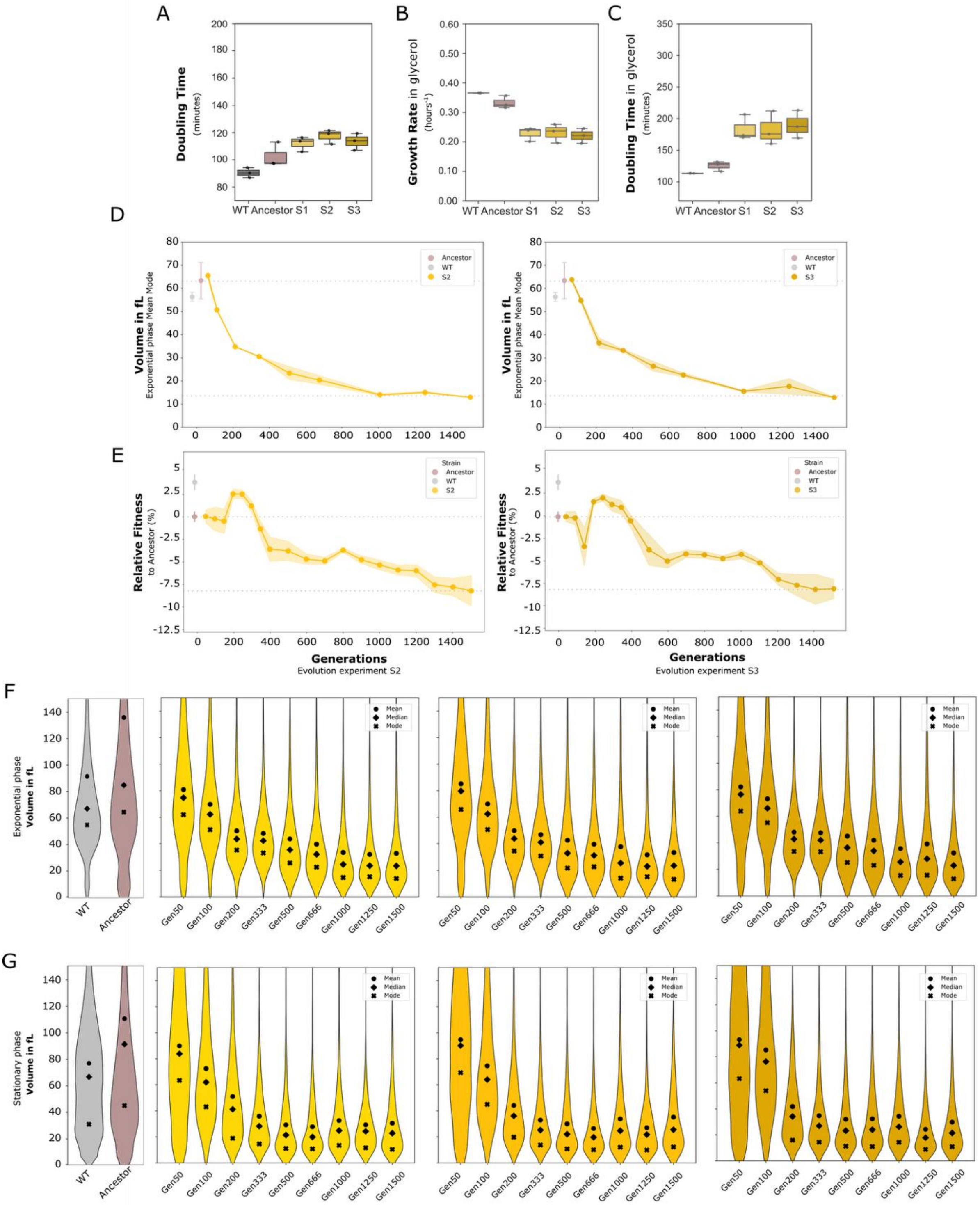

**Figure S3.** (A) Doubling times of reference and evolved strains (generation 1,500) in YPD. (B) Growth rates of reference and evolved strains in rich medium with glycerol as carbon source. (C) Doubling times of reference and evolved strains in glycerol medium. Box plots show medians, interquartile ranges, whiskers (1.5× IQR). (D) Size trajectories over 1,500 generations. Solid lines represent mean mode, and shaded areas indicate SD. (E) Trajectories of fitness relative to the ancestor over 1,500 generations. Solid lines represent mean fitness, and shaded areas indicate SD. (F–G) Cell size trajectories over 1,500 generations. Violin plots show average cell size distributions in exponential-phase (F) and stationary-phase (G). The first panel includes the WT and ancestor as references, followed by distributions for the three evolving populations across generations. Summary statistics highlighted were derived from all individual measurements.

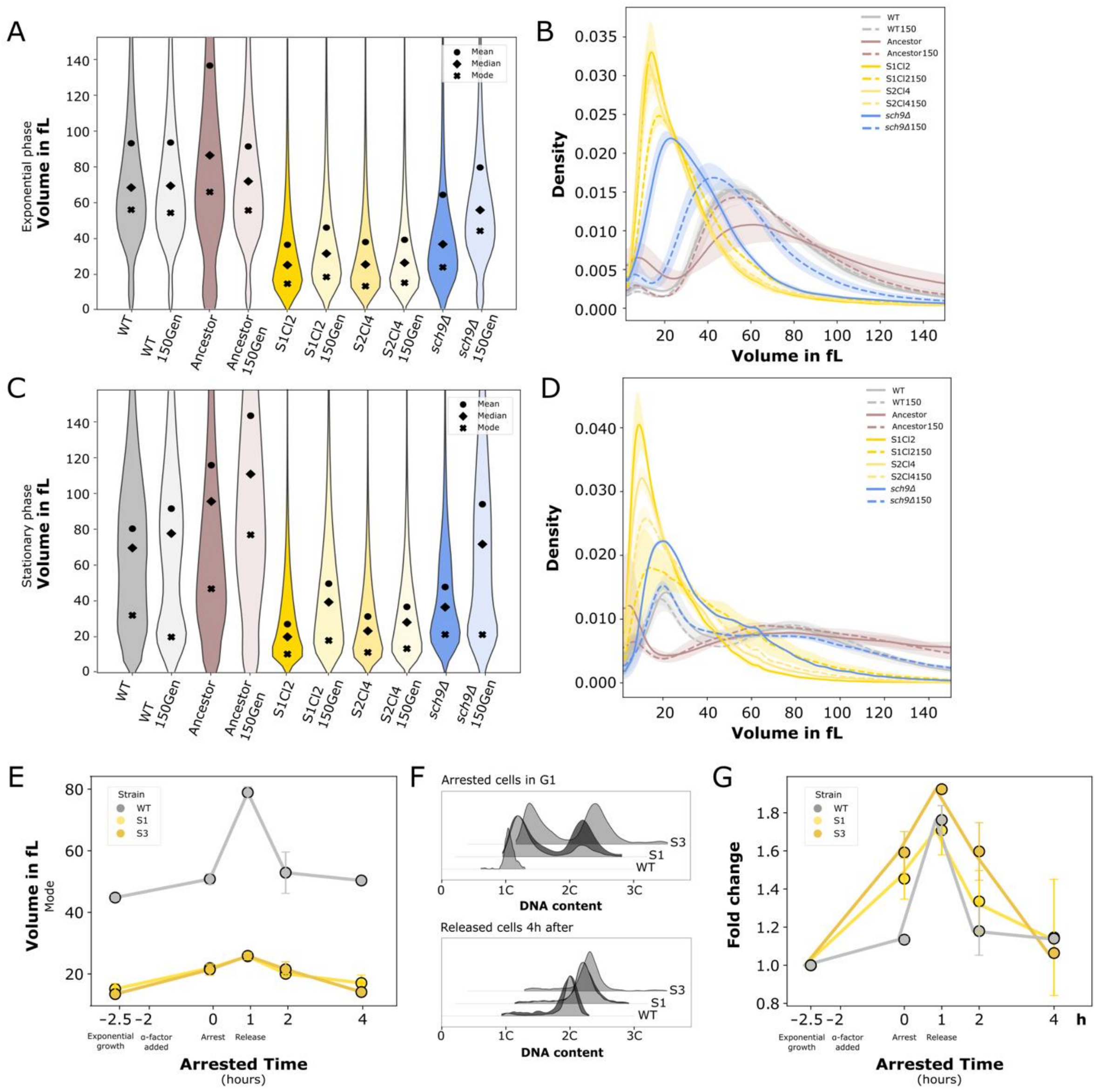


**Figure S4.** (A–D) Cell size distributions before and after 150 generations of evolution without size selection. Violin plots show average cell volumes (fL) in exponential-phase (A) and stationary-phase (C), while density plots show kernel density estimates of the same measurements in exponential-phase (B) and stationary-phase (D). For each strain, densities were computed per replicate and averaged to obtain a mean density curve, and shaded areas represent the standard deviation across replicates. Summary statistics highlighted were derived from all individual measurements. (E) Cell volume dynamics during synchronization with α-factor. Upon G1 arrest, cells maintained larger volumes, followed by a progressive decrease after release into fresh medium. (F) Cell cycle profiles from clones belonging to evolved pop S1 and S3: DNA content was quantified by SYTOX Green staining during exponential growth, α-factor arrest, and post-release. Arrested populations accumulated with a 1C DNA peak (G1), while released cells synchronously progressed through S phase and restored a 2C peak (G2/M). Evolved strains showed less efficient arrest, with a fraction remaining in G2/M. Genome sequencing revealed mutations in pheromone signaling genes, potentially explaining reduced responsiveness to α-factor. The x-axis shows arbitrary fluorescence units, the y-axis, relative frequencies. (G) Fold change in cell volume relative to the mean exponential volume before arrest.

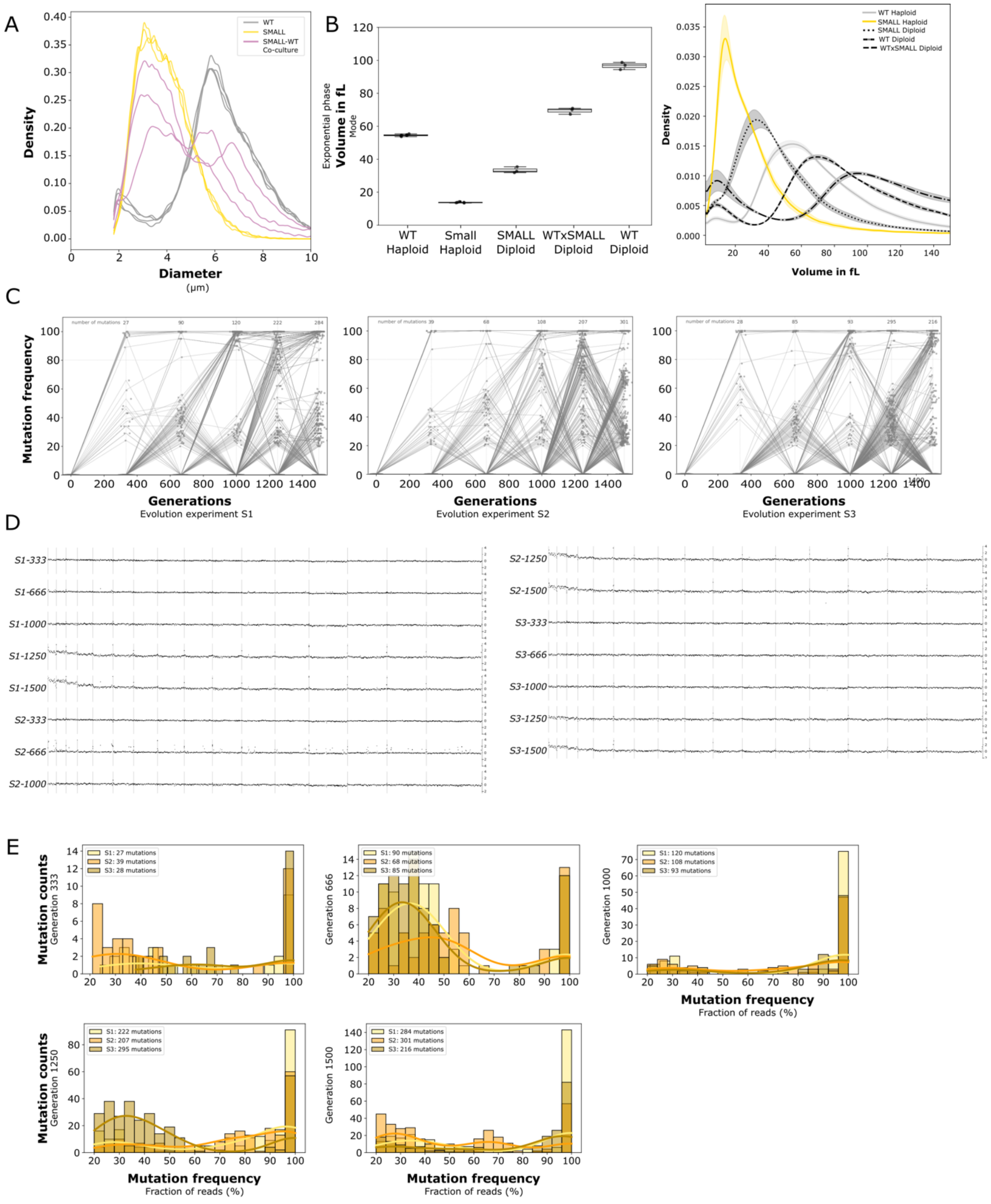


**Figure S5.** (A) Coexistence of WT and evolved Small-cell strains during 72 h co-culture. WT and Small strains were mixed 1:1 and grown together. Cell size distributions were measured using a Coulter Counter (n = 3). No significant shifts in population size distributions were observed. (B) Diploid cell size depends on parental strain combinations. Diploids were generated from crosses of Small × Small, Small × WT, and WT × WT strains. Cell volumes were measured for each diploid genotype. Small × WT diploids showed intermediate sizes, while WT × WT and Small × Small diploids retained the largest and smallest volumes, respectively, consistent with additive effects of parental cell size alleles. Density plots show cell distributions for diploids generated from crosses. For each strain, densities were computed per replicate and averaged to obtain a mean density curve. Shaded areas represent the standard deviation across replicates. (C) Allele frequency trajectories of mutated genes in Small populations S1–S3. (D) Copy number profiles of linear chromosomes across sequenced populations during experimental evolution (generations 333, 666, 1,000, 1,250 and 1,500). At 1,250 and 1,500 generations, apparent increases in small chromosome copy number are artifacts of DNA extraction bias, likely due to partial cell wall digestion that preferentially releases small chromosomes. (E) Distribution of mutated read fractions across Small populations. Mutation fraction (%) was calculated as the proportion of sequencing reads containing a given mutation. Histograms are overlaid with kernel density estimates (KDE, colored lines). Read fraction distributions were compared using Kolmogorov–Smirnov (KS) tests (full statistics in Supplementary file RawData-Figures).

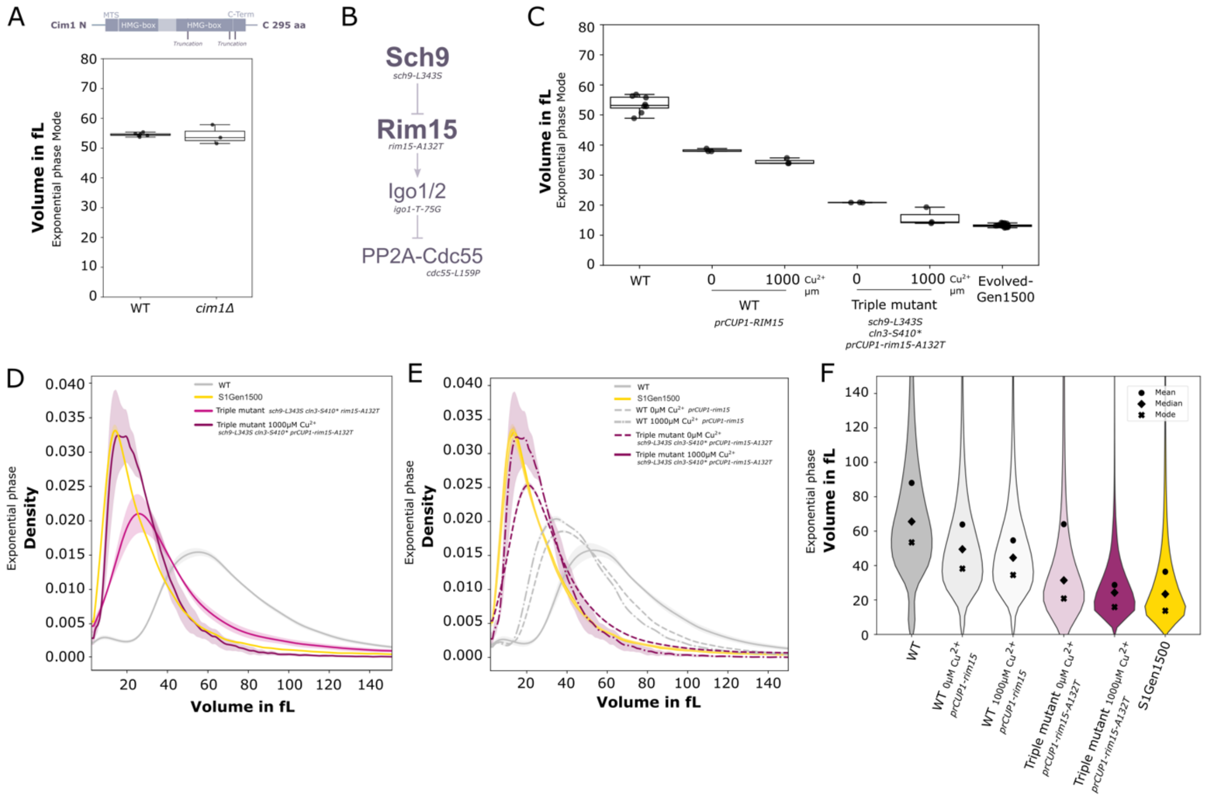


**Figure S6.** (A) Schematic representation of mutations identified in the *CIM1* gene, with affected domains indicated (adapted from Schrott and Osman (13)). Volumetric measurements (Coulter counter) of *CIM1* gene deletion. Box plots show medians, interquartile ranges, whiskers (1.5× IQR). (B) Schematic representation of the Greatwall kinase pathway with experimentally identified mutations. (C) Cell volumes measured in exponential-phase cultures (Coulter counter). Box plots show medians, interquartile ranges, whiskers (1.5× IQR). Response of WT and triple mutant strains to increasing copper concentrations (0 and 1000 µM $CuSO_4$) is shown; prCUP1 denotes expression of *RIM15* under control of the copper-inducible promoter. (D-E) Density distributions of cell volumes for the analyzed strains in exponential-phase. For each strain, densities were computed per replicate and averaged to obtain a mean density curve, and shaded areas represent the standard deviation across replicates. (F) Violin plots showing average cell distributions in exponential-phase. Summary statistics highlighted were derived from all individual measurements.

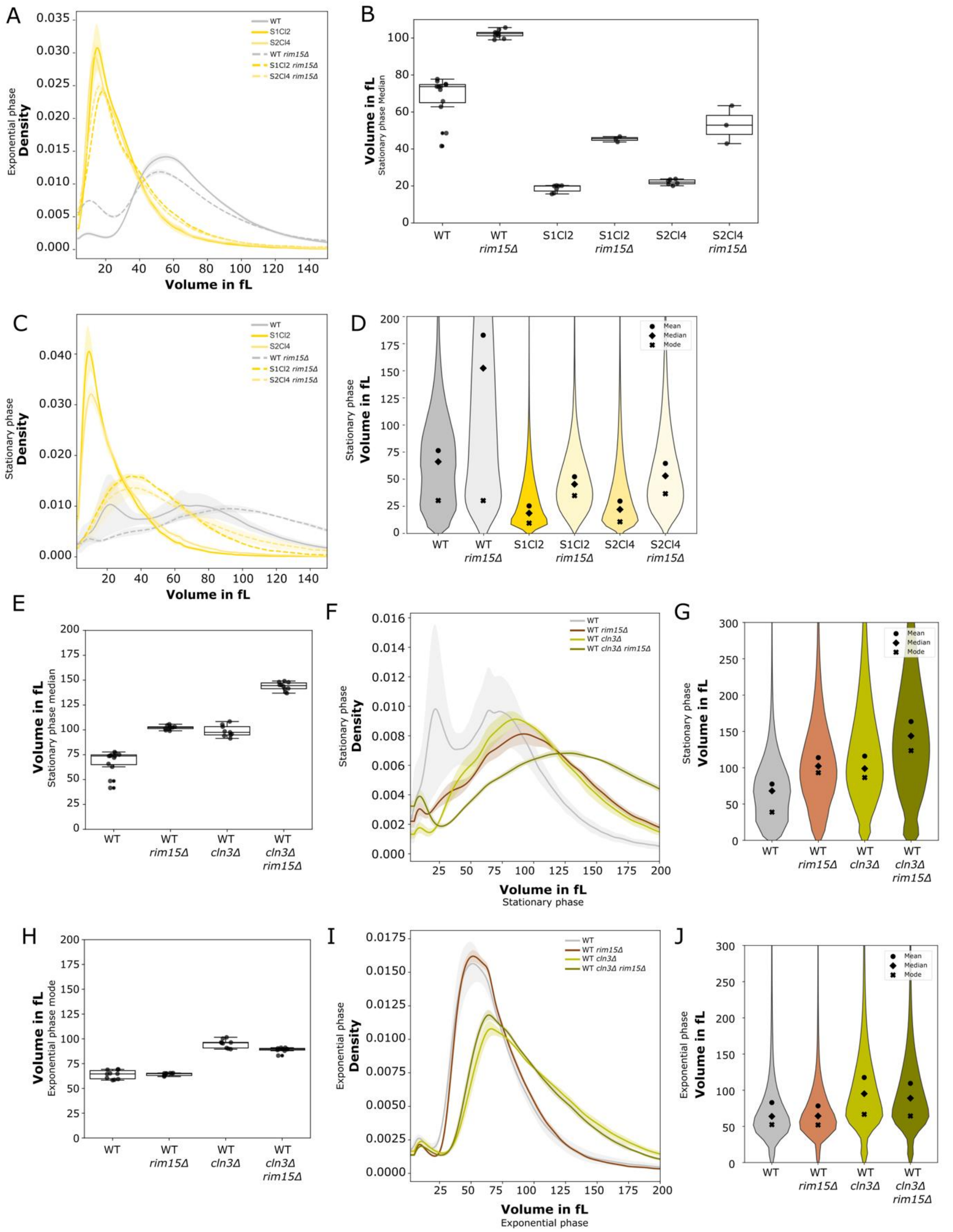


**Figure S7.** (A) Density plots showing cell size distributions of *RIM15* deletions in exponential-phase. (B) Median cell volumes and distributions for *RIM15* deletions in stationary-phase. (C) Average density distributions, and violin plots (D) for *RIM15* deletions in stationary-phase. (E-G)

Stationary-phase measurements for *rim15Δ*, *cln3Δ*, and *rim15Δ cln3Δ* double mutant: box plots of median cell volumes (E), density distributions (F), and violin plots (G). (H-J) Exponential-phase measurements for *rim15Δ*, *cln3Δ*, and *rim15Δ cln3Δ* double mutant: box plots of modal cell volumes (H), average density distributions (I), and violin plots (J). Samples were sonicated (5 × 1 s pulses at 10% amplitude) prior to measurement. Density distributions were computed per replicate and averaged to obtain a mean density curve. Shaded areas represent the standard deviation across replicates. Summary statistics highlighted in violin plots were derived from all individual measurements. Box plots show medians, interquartile ranges and whiskers (1.5 × IQR).

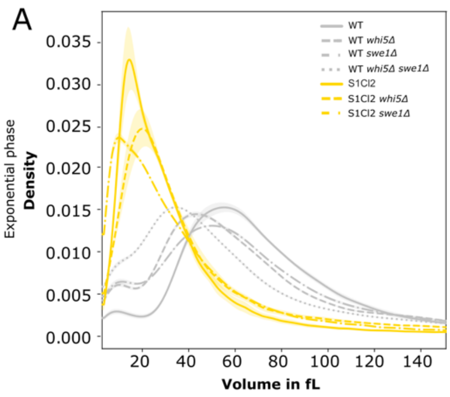


**Figure S8.** (A) Density distributions of cell volumes for *swe1Δ* and *whi5Δ* mutants in exponentially growing cultures. For each strain, densities were computed per replicate and averaged to obtain a mean density curve. Shaded areas represent the standard deviation across replicates.

### SUPPLEMENTARY TABLES

**Table S1.** Yeast strains used in this study.

List of *S. cerevisiae* strains employed in this work, including strain names, genotypes, and sources or construction details.

**Table S2.** Mutations identified in evolved lineages.

List of mutations identified in evolved yeast lineages. Each entry includes the gene name, SGD identifier, chromosome number, mutation type and position, and the subpopulation lineage in which the mutation occurred. Additional columns indicate the generation of detection, reference and mutant read counts, total reads, and the allele frequency (fraction).

**Table S3.** Genes previously implicated in cell size regulation.

List of genes identified in evolved yeast lineages that are reported in the literature to affect cell size in *S. cerevisiae*. The full list was obtained from the Saccharomyces Genome Database (SGD, https://www.yeastgenome.org/). Each entry provides the gene name, systematic name, associated phenotype, experimental approach, and experiment type category. Additional columns summarize mutant information, strain background, specific experimental details, and the corresponding reference.

**Table S4.** Genes under parallel evolution.

List of genes identified through statistical analysis as mutated more often than expected given their coding sequence length and the observed mutation rates, indicative of parallel evolution. Each entry includes the systematic name, SGD identifier, gene name, gene size, number of independent mutation hits, and counts of mutations in coding (CDS), noncoding (non-CDS) sequences, or synonymous mutations. Additional columns provide mutation frequency, zygosity, origin, gene description, and significance values (P and Bonferroni-corrected P (Pe)).

**Table S5.** Mutations identified by bulk segregant analysis.

List of mutations detected after two consecutive rounds of backcrossing and selection for small cell size, with corresponding SGD identifiers for each mutation.

**Table S6.** Summary of genes identified across analyses.

List of genes identified as putative adaptive candidates across the four complementary analyses (Tables S2–S5). Each entry includes the systematic name, counts and indicators used for ranking; SGD identifier, gene name, gene size, origin, and functional description.

**Supplementary Dataset S1.** RawData-Figures.

Excel file containing all raw numerical data underlying the figures presented in this study.
